# Size-dependent transition from steady contraction to waves in actomyosin networks with turnover

**DOI:** 10.1101/2022.07.13.499885

**Authors:** Ashwini Krishna, Mariya Savinov, Niv Ierushalmi, Alex Mogilner, Kinneret Keren

**Affiliations:** Department of Physics, Technion– Israel Institute of Technology, Haifa 32000, Israel; Courant Institute of Mathematical Sciences, New York University, New York, NY 10012, USA; Department of Biology, New York University, New York, NY 10012, USA; Network Biology Research Laboratories and Russell Berrie Nanotechnology Institute, Technion– Israel Institute of Technology, Haifa 32000, Israel

## Abstract

Actomyosin networks play essential roles in many cellular processes including intracellular transport, cell division, and cell motility, exhibiting a myriad of spatiotemporal patterns. Despite extensive research, how the interplay between network mechanics, turnover and geometry leads to these different patterns is not well understood. We focus on the size-dependent behavior of contracting actomyosin networks in the presence of turnover, using a reconstituted system based on cell extracts encapsulated in water-in-oil droplets. We find that the system can self-organize into different global contraction patterns, exhibiting persistent contractile flows in smaller droplets and periodic contractions in the form of waves or spirals in larger droplets. The transition between continuous and periodic contraction occurs at a characteristic length scale that is inversely dependent on the network contraction rate. These dynamics are recapitulated by a theoretical model, which considers the coexistence of different local density-dependent mechanical states with distinct rheological properties. The model shows how large-scale contractile behaviors emerge from the interplay between network percolation essential for long-range force transmission and rearrangements due to advection and turnover. Our findings thus demonstrate how varied contraction patterns can arise from the same microscopic constituents, without invoking specific biochemical regulation, merely by changing the system’s geometry.

## Introduction

Active networks composed of actin filaments and motor proteins are ubiquitous in cells, displaying a rich spectrum of dynamic behaviors (reviewed in ^1-3^). In general, actomyosin networks have a clear bias towards contraction ^4, 5^, that can be local or extend throughout the system with sustained or pulsatile temporal dynamics (e.g. ^6-10^). These diverse contractile patterns play a central role in driving the dynamic behaviors of living cells and tissues ^3, 5^, often exhibiting abrupt transitions along the cell cycle or during development (e.g. ^6-8^). Despite extensive research, how cells control and regulate their actomyosin cytoskeleton, generating networks with distinct architectures and contractile dynamics from the same underlying molecular components, is still not well understood. In particular, the respective roles of the intrinsic self-organized dynamics of the actomyosin cytoskeleton versus upstream regulation by various signaling pathways (e.g. Rho GTPases) in shaping these large-scale patterns often remains unclear ^11^.

*In vitro* realizations of systems containing actin filaments, crosslinkers and myosin motors have been instrumental for uncovering the properties of actomyosin networks and relating their large-scale dynamics to the underlying molecular composition ^12-16^. Most of the work on reconstituted actomyosin networks has focused on systems with limited turnover, in which contraction is typically a “one-shot” process ^12-16^. Extensive work has shown that the contractile behavior in these systems is primarily dependent on two effective parameters: the network connectivity and motor activity ^12, 13, 17, 18^. The structure and connectivity of the network are crucial for long-range transmission of the motor-generated forces, and hence dramatically influence the stress distribution within the network. At the same time, these forces generate flows and modulate the network structure. This feedback between force generation and network architecture can generate different large-scale behaviors including contracting states and quiescent states where the motor-generated forces are balanced by the rigidity of the network ^18^.

The introduction of continuous turnover, with rapid actin assembly and disassembly, in addition to motor activity, further complicates the interplay between network architecture and force generation, facilitating a richer spectrum of dynamic behaviors ^19-21^. Importantly, actomyosin networks *in vivo* are typically characterized by rapid turnover rates, which can be two orders of magnitude higher than the turnover rates of purified actin filaments, thanks to a host of auxiliary proteins involved in stimulating debranching, filament severing and depolymerization ^22^. Theoretical work indicates that the presence of continuous turnover makes contraction more sustainable ^19-21^, facilitating large-scale connectivity and force transmission even when the network is continuously flowing and rupturing. This can lead to a steady state of contraction ^23^, but under some conditions pulsatile contraction is also predicted ^19^. Despite their importance, our understanding of the behavior of actomyosin networks with rapid turnover is still rather limited. In particular, what determines the characteristic length scales for contraction (local vs global contraction) as well as its temporal dynamics (continuous vs pulsatile) in the presence of turnover is still not well understood.

Here we study the contractile behavior of bulk actomyosin networks with rapid turnover, using a reconstituted system based on cell extracts ^23-26^. Previous work has shown that bulk actomyosin networks can exhibit a steady state of contraction in the presence of turnover ^23^ or periodic contraction in the form of large-scale waves ^27-29^, but what determines the observed mode of contraction remained unclear. Our reconstituted system enables us to explore the behavior of actomyosin networks across a wide range of conditions and geometries, and show that these two apparently distinct modes of global contraction can arise from the same microscopic constituents and interactions, merely by changing the system’s size. We find that for different biochemical compositions, the system generically exhibits a transition from continuous contraction in smaller cells, to periodic contraction in the form of waves or spirals in larger cells. The characteristic length scale for this transition varies between the different conditions, and is found to increase in conditions exhibiting slower contraction.

The size-dependent transition from continuous to periodic contraction, which we observe under various conditions, was not predicted by any of the previous theoretical models or agent-based simulations ^19-21, 23, 30, 31^. To account for this phenomenology, we thus propose a new theoretical model that considers the interplay between network percolation required for long-range force transmission and advection due to the contractile flows driven by the motor-generated forces. Within this model, we take into account the heterogeneities that develop in the system due to network assembly, disassembly and advection, and consider the coexistence of different density-dependent mechanical states with distinct rheological properties in different regions of the system. The contractile behavior emerges from the self-organized dynamics of these coexisting local mechanical states. Importantly, the presence of rapid rates of actin turnover allows the system to efficiently bridge disjoint network regions, and generate global contraction even when contraction initiates locally. We show that our model accounts for the observed size-dependent transition in the global contraction patterns, with steady state continuous contraction predicted in smaller systems and periodic contraction developing when the system size exceeds a threshold value. Altogether, our experimental and theoretical results show how elaborate self-organized spatiotemporal patterns can emerge from the interplay between network structure, internal force generation and geometry, without necessitating additional regulation through specific signaling processes.

## Results

### Continuous and periodic contraction patterns in actin networks with rapid turnover

Contracting actin networks are generated in cell-like compartments by encapsulating cytoplasmic *Xenopus* egg extract in water-in-oil emulsion droplets ^23-26, 29, 32^. Due to the presence of rapid actin turnover rates (∼1 min^-1^) ^23, 25^, the system can sustain contracting network flows for hours. Interestingly, we observe two qualitatively different patterns of global contraction: (a) continuous contraction and (b) pulsatile, periodic contraction (Fig. 1, Supplementary Video 1). In the continuous regime, observed in smaller droplets, the system is essentially at steady state, as previously reported ^23^. In this case, the network exhibits a smooth density distribution that decreases towards the droplet’s periphery and a radial velocity profile in which the inward network flow speed increases linearly as a function of the distance from the contraction center (Fig. 1a; Supplementary Video 2). In the pulsatile regime, observed in larger droplets, the system exhibits periodic variation in the form of concentric waves in both network density and flow speed (Fig. 1b; Supplementary Video 3) ^27-29^. Wave fronts of higher actin density appear regularly near the periphery of the droplet, and contract inward toward the contraction center (where an aggregate of cellular debris forms, whose size scales with the size of the droplet ^23, 29^). The shape of the contracting wave fronts reflects the shape of the boundaries of the system ^28^; e.g. circular wave fronts form in round droplets (Fig. 1b; Supplementary Video 3), whereas in elongated channels the wave fronts are straight, parallel to the channel walls (Fig. S1; Supplementary Video 4). The pulsatile contraction has a characteristic time period of ∼1min, that is evident in the radial kymographs of the network density and velocity (Fig. 1c). We observe qualitatively similar behavior in different extract batches, but quantitatively there is some variation (e.g. in wave period and the contraction rate) between different extracts and also depending on the extract concentration used (60-100%; see Methods).

**Figure 1.**
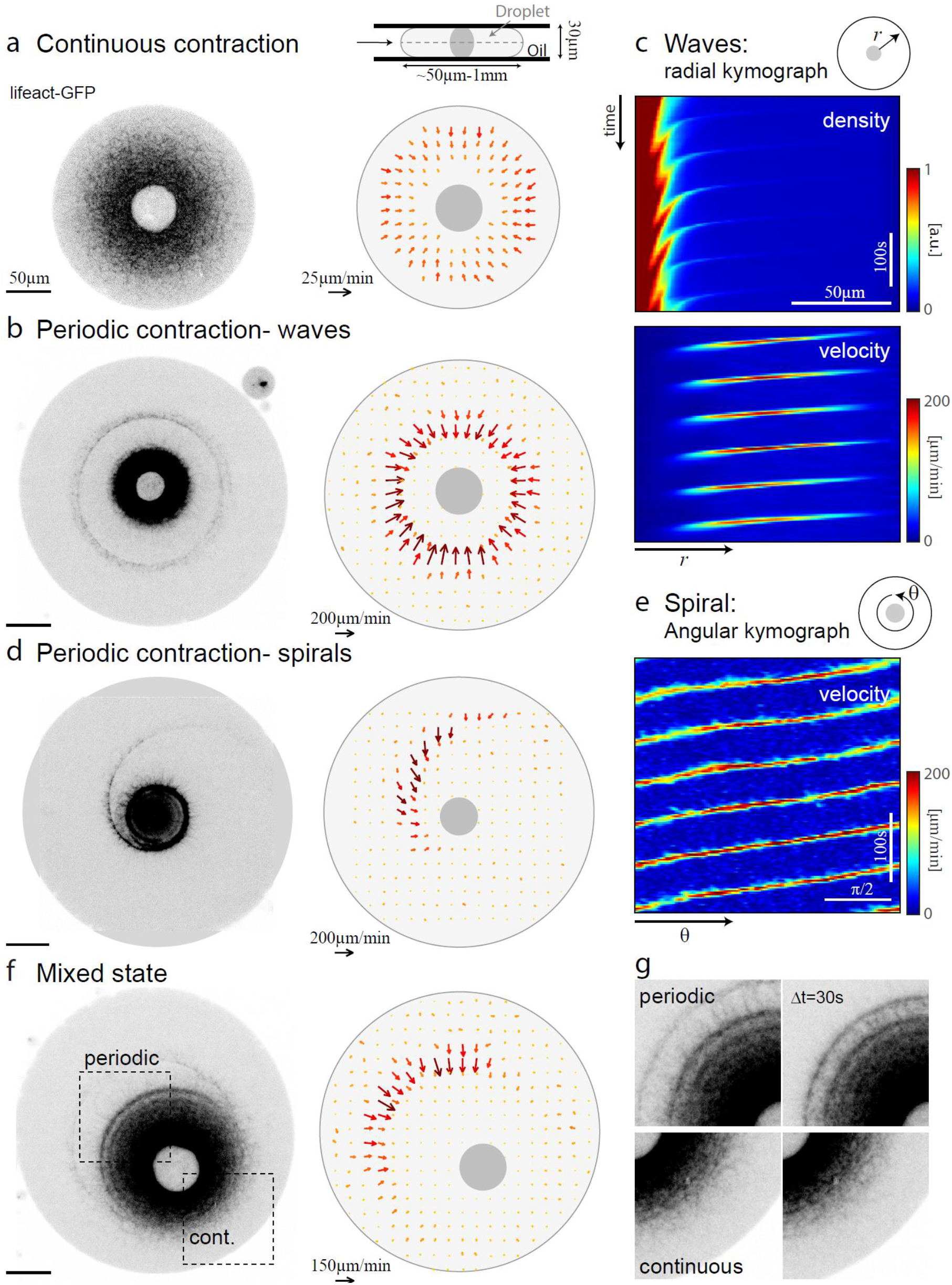
Examples of continuous and periodic contraction patterns in actin networks with rapid turnover. (a,b) Spinning disk confocal images (left; inverted contrast) and the corresponding velocity fields extracted using optical flow (right; see Methods) of water-in-oil droplets exhibiting continuous contraction (a; Supplementary Video 2) or periodic contraction in the form of concentric waves (b; Supplementary Video 3). The droplets are sandwiched between two coverslips in a pancake-like geometry, and the images are taken at the droplet’s mid-plane (a-inset; Methods). (c) Radial kymographs showing the periodic variation in density (top) and network flow velocity (bottom) over time in the droplet shown in (b). (d) Spinning disk confocal images (left) and the corresponding velocity fields (right) of a water-in-oil droplet exhibiting periodic contraction in the form of a clockwise spiral (Supplementary Video 5). (e) Kymograph showing the angular variation in velocity over time for the spiral shown in (d). The linearly-varying phase of the spiral wave front as a function of angle generates periodically-spaced parallel diagonal lines in the angular kymograph. (f) Spinning disk confocal images (left) and the corresponding velocity fields (right) of a water-in-oil droplet in which the contraction center is positioned at an off-centered location. The actin network exhibits different contractile behaviors in different regions of the droplet (Supplementary Video 6). (g) Subsequent images from different regions of the droplet shown in (f) depicting the contraction of a wave front on the far side of the droplet (upper left box in (f)) and continuous contraction on the opposite side (lower right box in (f)). The droplets shown contain 95-98% *Xenopus* cell extract supplemented with lifeact-GFP to visualize the actin network (Methods).

Interestingly, the periodic contraction can also take the form of spirals (Fig. 1d, Supplementary Video 5). In this case, the local network density and velocity exhibit periodic modulations as in the circular concentric waves, but the wave front forms a spiral. The contractile flow is primarily directed inward, but there is also a non-zero tangential component along the spiral wave front (Fig. 1d). We observe spirals rotating both clockwise and counterclockwise with similar probabilities (Fig. S2), suggesting that the chiral patterns arise from spontaneous angular symmetry breaking. The spirals can be stable for multiple rotations (Fig. 1e), but occasionally we observe transitions between concentric waves and spirals (Fig. S3). The transition into a spiral is typically preceded by angular variation in the network density distribution that persists at the same location over multiple consecutive wave periods (Fig. S3b). These spontaneous angular variations appear sufficient for driving the transition between the two types of periodic contraction patterns, namely concentric waves and spirals.

The continuous and periodic contraction regimes can sometimes be observed together within a single droplet in which the contraction center is located asymmetrically (Fig. 1f, Supplementary Video 6). In such cases, the periodic contraction occurs on the side of the droplet where the distance between the contraction center and the droplet’s periphery reaches a maximal value, whereas continuous contraction is apparent on the opposite side (Fig. 1g). These observations indicate that the pulsatile contractions in larger droplets are not caused by scaling of biochemical concentrations with the droplet’s size, but rather that the transition between continuous and periodic contraction behavior directly depends on the system’s geometry, specifically on the distance between the boundary of the system and the contraction center.

### Transition from continuous to periodic contraction depends on system size and contraction rate

To characterize the size-dependent transition between continuous and periodic contraction, we observe droplets of different sizes (naturally formed in the emulsion-generating process) and assess their global contractile behavior by looking for periodic density modulations in the radial kymographs (Fig. 2a; Methods). The properties of the reconstituted actomyosin networks can be modulated by adding different components of the actin machinery or pharmacological drugs that modify their activity ^23^. Interestingly, while the network appearance and properties can change substantially, we observe a qualitatively similar transition from continuous contraction in smaller droplets to periodic contraction in larger droplets under a range of conditions (Fig. 2, Supplementary Video 7). The characteristic length scale for the transition from continuous to periodic contraction, however, shifts depending on the system’s composition (Fig. 2b,c). All comparative analysis between different conditions was done with the same batch of extract, but qualitatively the observed trends were similar in different extracts. For our standard conditions (80% extract), the characteristic length scale for the transition between continuous and periodic contraction is ∼150 μm (Methods). Enhancing actin assembly by adding nucleation-promoting factors, such as mDia1, which nucleates unbranched actin filaments, or ActA, which activates the Arp2/3 complex to nucleate branched actin filaments, decreases the contraction rate and shifts the transition length to larger values (Fig. 2b-d). Conversely, adding Capping Protein which caps growing actin filaments increases the contraction rate and shifts the transition length to smaller values. Within the periodic regime, the wave period varies between different conditions (Fig. 2e), but does not have a strong dependence on system size (Figs. 2f, S4; see also ^29^).

**Figure 2.**
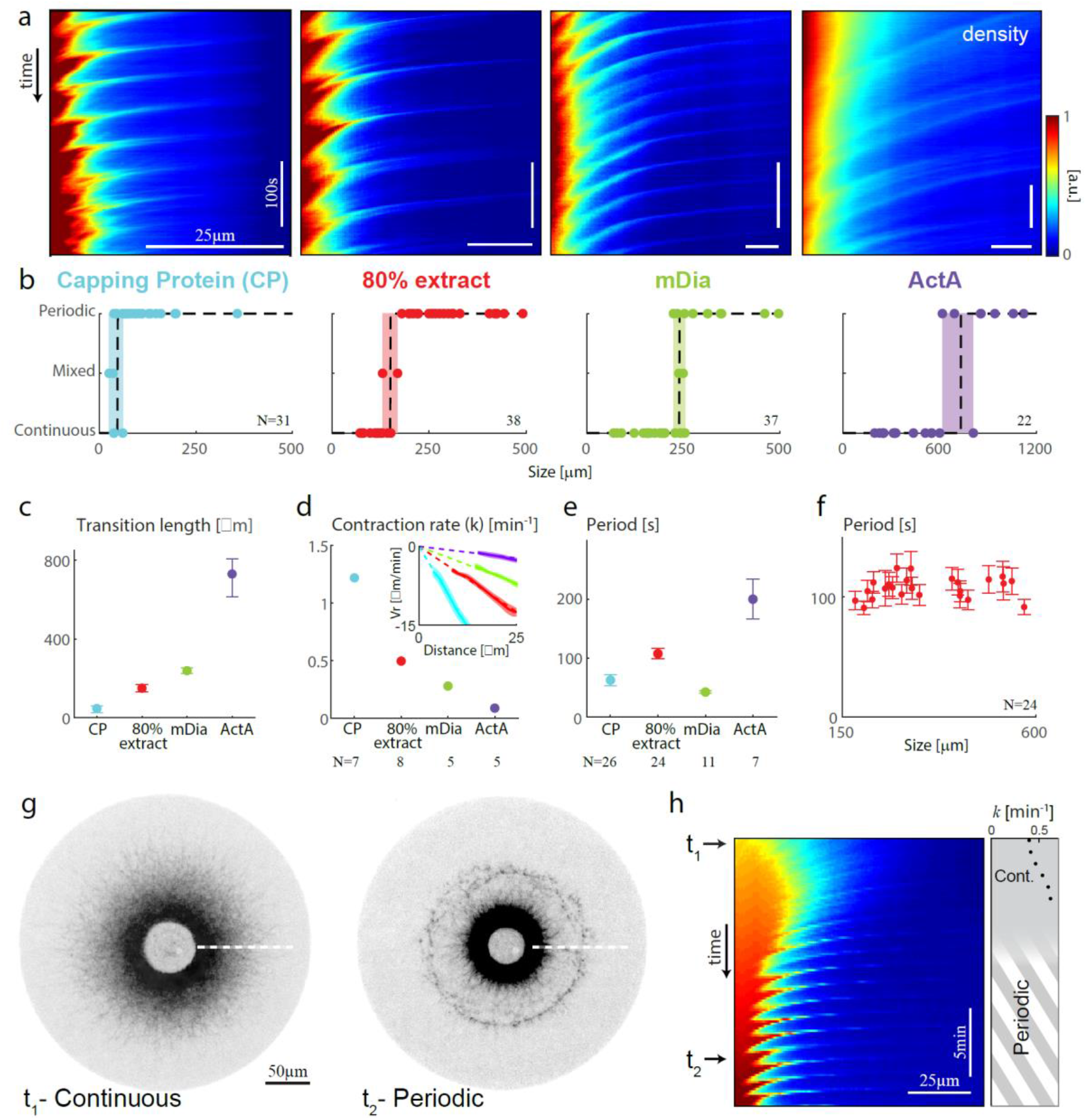
Transition from continuous to periodic contraction depends on system size and contraction rate. (a) Radial kymographs showing the periodic variation in network density over time for droplets with different compositions (Supplementary Video 7). The droplets contain 80% *Xenopus* cell extract supplemented with lifeact-GFP to visualize the actin network and auxiliary proteins as indicated (1μM Capping protein, none, 0.5μM mDia, or 1.5μM ActA; see Methods). (b) The global contractile behavior of droplets with different compositions (as in a) was determined as a function of the distance between the contraction center and the droplets’ boundary. For each condition, large droplets exhibit periodic contraction and small droplets exhibit continuous contraction. The transition length for each condition varies as indicated (dashed line). The shaded region corresponds to the size region between the smallest droplet exhibiting periodic behavior and the largest droplet showing continuous contraction. (c) The transition length estimated from the data shown in (b) for the different conditions (error bar corresponds to the width of the shaded region in (b)). (d) The contraction rate in the continuous regime for the different conditions. The contraction rate was measured in small droplets (R< 70μm) from the slope of the radial velocity as a function of distance from the contraction center ^23^ (Inset; Methods). (e) The wave period in the periodic regime for the different conditions. For each condition, the mean±std over different droplets is shown. (f) The wave period for droplets with 80% *Xenopus* cell extract as a function of system size. The error bar corresponds to the uncertainty in determining the period in each droplet (Methods). (g) Spinning disk confocal images of a water-in-oil exhibiting a transition from continuous contraction (left) to periodic contraction (right). The droplet contains 98% *Xenopus* cell extract and is imaged over 25min, during which the contractile behavior of the droplets changes gradually (Supplementary Video 8). (h) Radial kymograph showing the variation in density over time for the droplet shown in (g). The arrows indicate the time points corresponding to the images shown in (g). Right: the increasing contraction rate in the continuous regime is plotted as a function of time. The appearance of periodic modulations in the density is indicated.

We have previously shown that in the continuous contracting regime, the network undergoes telescopic contraction with a nearly homogenous, density-independent contraction ^23^. As such, the contraction of these spatially inhomogeneous networks can be characterized by a single characteristic contraction rate. While this contraction rate is an emerging property of the system that depends in a non-trivial manner on the internal force generation and architecture of the network, as well as on the system size (Fig. S5), it can nonetheless be measured in a straightforward manner from the slope of the radial velocity profile (Fig. 2d inset). Interestingly, we find that for the different conditions examined, the transition length is anti-correlated with the contraction rate (Fig. 2c,d). The characteristic time period in the wavy regime also varies between different conditions, but does not appear to be correlated with the contraction rate (Fig 2d,e). Notably, the wave period is of the same order of magnitude as the duration of the actin turnover cycle in the system (∼1min ^23^). Overall, these observations indicate that while the quantitative characteristics of the transition from continuous to periodic contraction vary depending on system properties, the appearance of such a transition is a generic, self-organized feature of contracting actin networks with turnover, that does not depend on fine-tuning of system parameters.

The nature of the transition between continuous and periodic contraction can be visualized by following individual droplets over extended time scales. The extract is a complex mixture of components whose properties can gradually change over time. We typically observe a slow upward drift in the contraction rate over time that is more pronounced in some extracts. While the source of this time-dependent behavior is not well-characterized, we can nevertheless harness this gradual variation in system properties to follow the transition between continuous and periodic contraction and characterize the onset of periodic contraction in a single droplet (Fig. 2g,h, Supplementary Video 8). As the contraction rate increases over time (Fig. 2h, right), the characteristic transition length to the wavy regime decreases, in agreement with our observations with varying system composition (Fig. 2b). As a result, individual droplets that are initially just below the transition length and exhibit continuous contraction, change their global contractile behavior over time and transition into the periodic regime (Fig. 2g,h). Over time, the continuous network flow first breaks into short intermittent arcs that are not well-synchronized in space and time. These local modulations become more spatially synchronized, turning into well-defined wave fronts that increase in amplitude as the system moves further away from the transition point.

### Modeling the contractile behavior of actomyosin networks with turnover

To understand the observed contractile behaviors, and in particular how the system can switch from continuous contraction to periodic waves, we turn to theoretical modeling. The simplest description of a dynamic contractile actomyosin network introduced in ^23^, posits that the network assembles and disassembles, behaving effectively as a highly viscous fluid on long time scales, with an active contractile (myosin-powered) stress that drives network contraction against its internal viscous resistance. While this simple model can account for the continuous steady-state contraction observed in smaller droplets ^23^, it cannot generate periodic contractions.

An obvious limitation of this simple formulation is that both the contractile stress and the effective network viscosity are assumed to be finite for any density. Previous work has shown that contraction in actomyosin networks occurs only above a minimal threshold for network connectivity and motor activity ^12, 13^, implying that the network behavior at low densities must be qualitatively different. Theoretically, this transition has been described in terms of percolation models, that consider network connectivity and its interplay with motor activity (reviewed in ^18^): the network must be connected (percolated) for myosin motors to generate long-range contractile forces and at the same time motor activity modulates network structure. In the presence of continuous actin turnover, this picture must be further modified to include the effects of network disassembly and reassembly and advection generated by persistent network flows.

Here we introduce density dependencies of the contractility and viscosity into the active fluid model and consider the coexistence of different local mechanical regimes. At low filament density, the network will not be percolated and hence we assume its effective viscosity will be negligible and force propagation in space is limited. Taking this into account, qualitatively, waves could arise via cycles of contraction and gelation as previously suggested ^27, 28^: after a contractile wave sweeps the network inward, unconnected filaments and network fragments reassemble at the droplet’s periphery. Only after a finite time interval, a new wave is triggered when the local density increases beyond the contraction threshold. Note, that to generate periodic global waves in this scenario, one must assume that there is also an intermediate regime where the network is percolated but not yet contractile. Consider the wake of a contracting front. As the unconnected network reassembles, the density will first reach the contraction threshold near the periphery, furthest from the previous wave front (since the network there had the longest time to reassemble). When this happens, the nascent network in the middle (between the center and the periphery) must be percolated, to provide a mechanical connection between the network near the boundary and the previous wave front. Otherwise, the system will form multiple locally contracting clusters rather than global periodic contraction waves (see Supplementary Information (SI)).

This cyclic gelation-contraction mechanism also provides an intuitive explanation for the observed size-dependence of the transition from continuous to periodic contraction. Since the contractile network elements are connected in series, their contractions add up across the droplet, so the contractile flow at the periphery increases with droplet size. In smaller droplets, the inward flow at the periphery is slow enough for assembly to replenish the receding network continuously. In larger droplets, however, the inward flow wipes out the network so rapidly that a finite period of reassembly (without flow) is necessary before another cycle of contraction can initiate.

To test this picture, one needs simulations of a quantitative model. We base our model on the previously suggested state-diagram for actomyosin networks (reviewed in ^18^), with some notable differences (Fig. 3a). That state diagram ^18^, depicted as a function of two parameters – network connectivity (an effective measure that depends on network density, filament lengths, as well as the concentration and properties of crosslinking proteins) and motor activity, includes four states: an unconnected “gas”, a percolated but non-contracting network, a globally contracting network, and a locally contracting network. It is reasonable to assume that the connectivity is an increasing function of actin density, while the motor activity is an increasing function of myosin density. We thus employ a simplified state diagram, depending on a single network density field (assuming that both actin and myosin densities are proportional to this network density), and consider three density-dependent mechanical regimes (Fig. 3a), namely, an unconnected network fragments (“gas-like”) state (I), a connected (but not contracting) state (II), and a contracting state (III). The fourth local-contraction regime emerges as a small-scale instability in our model (see below).

**Figure 3.**
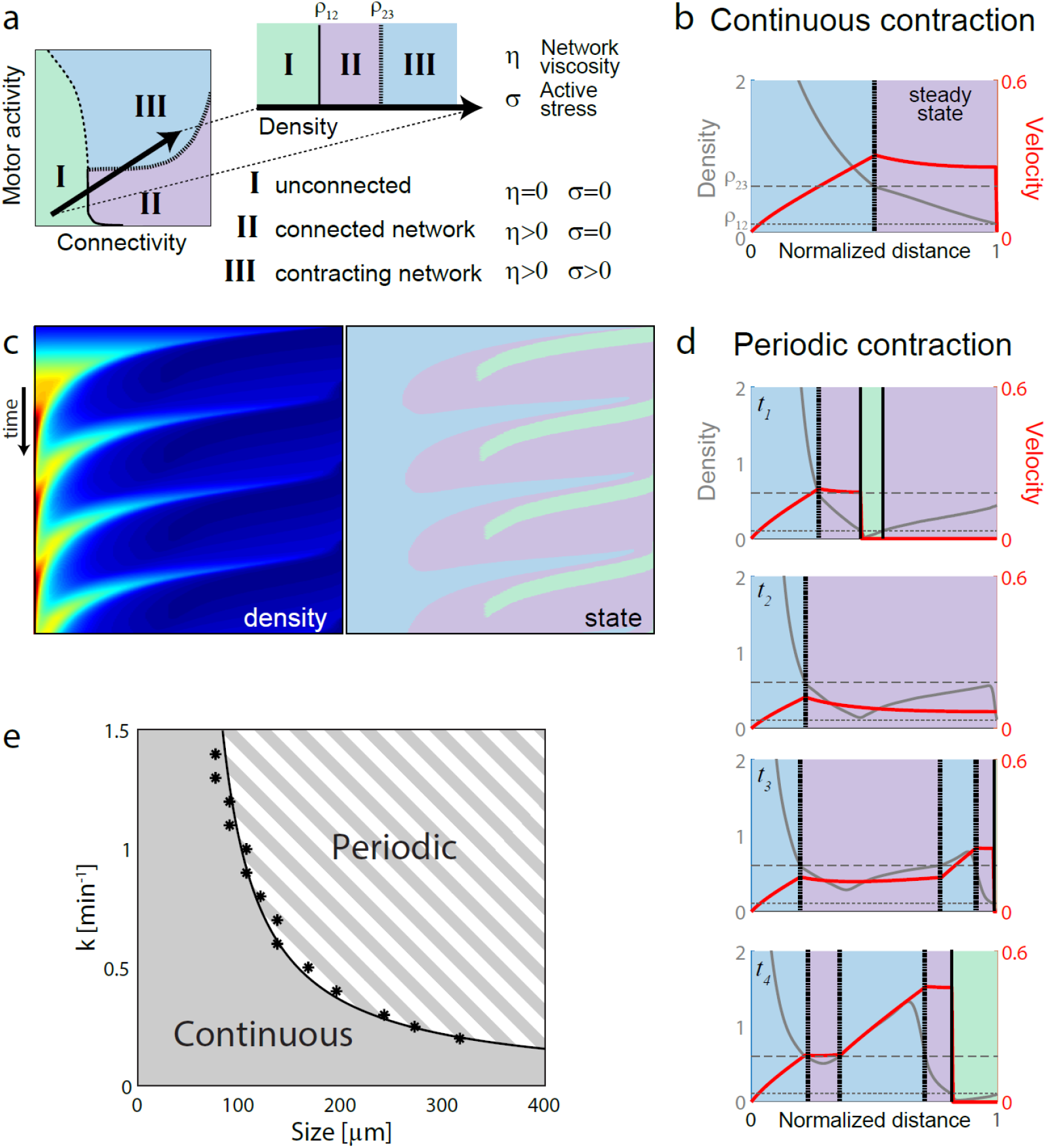
Modeling the transition between continuous and periodic contraction. (a) Schematic state diagram of the local contractile state as a function of network connectivity and motor activity adapted from ^18^ (left), and the simplified state diagram as a function of network density used in our model (right). (b) Graph showing the simulated steady state density (grey) and velocity (red) distributions as a function of distance from the contraction center determined from 2D simulations in the continuous contraction regime (the results are shown for the non-dimensionalized variables; see SI). In b,d the dotted and dashed horizontal lines indicate the percolation and contraction density thresholds, respectively. The colors demarcate the regions with unconnected (green), percolated (violet) and contractile (blue) network densities; the vertical dotted lines show the percolated-contractile region boundaries and vertical solid lines show the unconnected-percolation region boundaries. (c) Kymographs showing the periodic modulation of network density (left) and the corresponding local network state (right) along a radial cross section as a function of time, determined from 2D simulations. (d) Graphs showing the density (grey) and velocity (red) distributions at different time points during a wave cycle in the periodic contraction regime. The background colors in b,d indicate the local state of the network at that location. (e) State diagram of the global contractile behavior of the system as a function of the contraction rate (*k*) and the system’s size (*R*). The system behavior was simulated for different values of *k* and *R*. For each value of *k*, the transition radius for which periodic behavior is first observed is indicated (asterisks). The line separating the continuous and periodic contraction regimes is determined from a best fit to the transition length for different values of the contraction rate (*R*_*tr*_ ≈1.8*v*_0_ / *k* +1.5*v*_0_ / *β*).

Importantly, the state diagram in ^18^ referred to networks with negligible turnover, considering a primarily elastic network and irreversible contraction from an initial state. The presence of network turnover, induces qualitative changes, allowing connected network clusters to dynamically form and break and implying that the network must fluidize at long time scales (compared to the turnover timescale). Moreover, the different mechanical regimes can coexist in different regions of the system and the boundaries between these regions can be dynamic over time. We thus rely on the different mechanical states identified previously, but, in contrast to previous work, we assume the network to be primarily viscous on the long time scales considered (we argue in the SI that considering a purely elastic 2D or 3D network is incompatible with our data). Most importantly, we examine how the three mechanical regimes play out dynamically in space and time in the presence of continuous network turnover and advection.

Our model is based on the conceptually simple hydrodynamic model of actomyosin networks introduced in ^23^, that invokes (1) mass conservation in the presence of network turnover and (2) force balance. We consider a network with density *ρ* flowing with a velocity 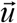. Mass conservation implies that the network density, *ρ*, changes due to assembly with rate*α*, disassembly with rate *β*, and drift with velocity 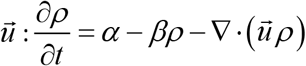. We use the force balance equation: 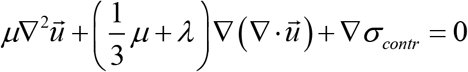, for which we assume that the total internal stress in the network is the sum of an active isotropic contractile stress,*σ*_*contr*_, and a passive viscous stress of an isotropic compressible Newtonian viscous fluid. In the force balance equation, the first two terms account for the divergence of this viscous stress, where *μ* and *λ* are the effective shear and bulk viscosities of the actin network. The friction due to movement of the actin mesh relative to the surrounding fluid is negligible ^32^. We explore this model in two dimensions (2D), to approximate the pancake-like geometry of our experimental system, and in one dimension (1D), to approximate the capillary geometry and to gain analytical insight (see additional discussion of the mechanics of the network in the SI).

We describe the different mechanical regimes of the network at three density ranges by defining the local network viscosity and active stress (Fig. 3a): (I) At low density, the network is disconnected, consisting of a “gas-like” solution of unconnected filaments and small network fragments diffusing in the solute, so we assume negligible viscosity and no active force generation. (II) At intermediate densities, the network crosses a percolation transition and becomes interconnected, but not yet contracting. In this regime the network resists deformations, so we assume its viscosity is finite, but the contractile stress is negligible (Two notes: 1) in the SI, we consider the possibility of a solid-like (elastic), rather than a fluid-like (viscous), interconnected network, and explain that the predicted behavior then is essentially the same in 1D and very different in 2D or 3D; 2) the likely reason for the network to be non-contracting in this regime is that the active myosin stresses are balanced by elastic stresses (generating a pre-stressed state); this is simplified here by ignoring the elastic stress and assuming negligible active stress). (III) At higher density, the network is both interconnected and contracting. Mathematically, we characterize the different regimes as follows: *μ* = *λ* = 0 if *ρ* < *ρ*_12_, *μ, λ* > 0 if *ρ* > *ρ*_12_ ; *σ*_*contr*_ = 0if *ρ* < *ρ*_23_, and*σ*_*contr*_ = *σ*_0_ > 0 if *ρ* > *ρ*_23_, where *ρ*_12_ and *ρ*_23_ are the critical densities for network percolation and contractility, respectively. Note that *ρ*_12_ < *ρ*_23_, since the network must be percolated to contract. The contractile regime can be characterized by a contraction rate *k*, which is the ratio of the active stress and an effective viscosity,*η*_0_, *k* = *σ*_0_ /*η*_0_, where*η*_0_ is a function of *μ, λ* and a geometric parameter (see SI). This rate, which has dimensions of inverse time, determines the spatial gradient of the contractile velocity and inverse characteristic time of the network contraction. In general, the viscosities and contractile stress are functions of the density within regions II and III of the state diagram, but for simplicity we approximate them as piece-wise-constant functions of the density. The contraction rate can also be density-dependent, but because of this approximation and based on our observations (Fig. 2d), we take the contraction rate to be a density-independent constant _23_.

To account for the boundary conditions in the system, we assume no flow at the dense innermost boundary, where the network sticks to the aggregate at the droplet’s center ^23, 32^. We further assume that actin filaments at the outer boundary of the interconnected network polymerize with an effective rate *v*_0_, so the free boundary of the network grows outward normally to the boundary with rate 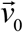. Thus, if the mechanical forces move the material points near this boundary with velocity 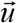, the network boundary moves with a net velocity 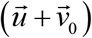. The parameter *v*_0_ reflects the rate of elongation of actin filaments at the network boundary, and is thus proportional to the actin assembly rate*α* :*α* ≈ *nv*_0_, where *n* is the local number of growing filaments per unit length (in 1D) or area (in 2D), providing the density is measured in units of filament length per unit length (1D) or area (2D), respectively.

We solve the model equations numerically (details in the SI). In the simulations, we take experimentally measured values or estimate realistic values for model parameters, such as polymerization rate *v*_0_ and network disassembly rate *β*, and vary two key parameters – the radius of the droplet, *R*, and the contraction rate, *k*. The model makes a simple prediction: if the droplet size is smaller than a critical size, *R* < *R*_*tr*_ = *cv*_0_ / *k*, where *c* is a dimensionless parameter of order unity, then a continuous steady centripetal network flow is maintained (Fig. 3b,e), as described previously ^23^. However, if the droplet radius exceeds the critical transition length, *R*_*tr*_, then pulsatile contraction waves emerge (Fig. 3c,e). Snapshots from the simulation (Fig. 3d, Supplementary Video 9), illustrate the key events in a wave cycle: (i) the contracting network recedes away from the droplets’ boundary until the periphery of the interconnected network stabilizes and remains stationary at some position *r*_12_ where *ρ* (*r*_12_) = *ρ*_12_ and 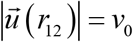 (so that 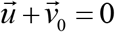). During this time the unconnected network density increases at the periphery; (ii) the connectivity threshold density is reached everywhere, and a new wave starts as a mechanical bridge is formed between the contracting network near the core and the newly assembled region extending to the droplet’s periphery; (iii) the density reaches the contractile regime in a region near the periphery, accelerating the wave; (iv) inward contraction of the network results in a region that is transiently unconnected near the droplet’s boundary. This model can also be used to simulate the dynamics of the system when the contraction rate is gradually increased over time, whereby a droplet of a given size can display a transition from continuous to periodic contraction (Fig. S6, Supplementary Video 10), as observed experimentally (Fig. 2g,h).

The global contractile behavior of the system can thus be characterized as a function of two important factors: the system size, *R*, and the contraction rate *k*, predicting a state diagram for steady/periodic contractile behavior in the *k* − *R* parameter space (Fig. 3e). The transition length (*R*_*tr*_) is predicted to be inversely proportional to the contraction rate as observed experimentally (Fig. 2c,d), which is easy to understand as follows. In the continuously contracting case, the interconnected network boundary is stationary, and hence flowing inward with a velocity equal to the growth rate of the network boundary, *v*_0_. Near the transition length, the radial position of the contracting network boundary is approximately *v*_0_ / *k*, while the radial position of the outward interconnected network boundary, which is a little farther outwards, is *cv*_0_ / *k* where *c* ∼ 1. The outward interconnected network boundary must be near the droplet’s interface, otherwise the unconnected fragments at the droplet periphery will periodically assemble and generate periodic contractions. Thus, *R*_*tr*_ ∼ *cv*_0_ / *k* (see detailed analysis in SI). This argument further suggests that the wave period should be determined by the time interval required for the network to assemble from scratch at the periphery up to the critical density, which is of the order of the network turnover timescale, or the inverse disassembly rate *β* (see SI). Indeed, the simulations show that the wave period is a few-fold 1/ *β*, and is largely insensitive to system size *R* and the contraction rate *k* (Fig. S7), as observed experimentally (Figs. 2f, S4).

The importance of the assumption that the interconnected network edge grows due to actin polymerization becomes clear: without it, the contractile network would recede at ever slowing, but finite speed toward the center, leaving a small gap with an unconnected region just behind it. This gap would never allow the growing density at the periphery to connect to the central contracting core, generating contraction in opposite directions, toward the center and toward the periphery. Alternative mechanisms that could induce global contraction are also possible. One possibility is that there are long non-contractile actin bundles permeating the active network and creating a non-stretchable but easily compressible, cable net, which connects the actively contractile local domains of the actomyosin network ensuring global contractions. We discuss this and other alternative models in the Discussion section and in the SI.

### Transition from global to local contraction with rapid network turnover

Previous work identified a transition from global contraction that span the system’s size to local contraction, where the network breaks down into multiple discrete clusters ^12, 15, 16, 18, 19^. This transition was shown to occurs at high motor activity accompanied by limited network connectivity. A discrete agent-based model indicates that in the presence of turnover a similar transition occurs (SI). Notably, the periodic regime appears as an intermediate between global continuous contraction and local contraction; when the network connectivity diminishes, the periodic contraction regime breaks down into a more irregular contractile behavior (SI, Fig. S8). In particular, contractile ‘asters’ evolve: local high-density clusters characterized by small size and rapid centripetal flow, appearing at random locations, with large spaces between them that are devoid of an interconnected network (Fig. S8). Interestingly, we also find an intermediate regime, where the contraction pattern alternates repeatedly between local contraction into clusters that subsequently get connected at random times with a dense central steadily contracting core cluster (Fig. S8). This intermediate regime, that exhibits contraction on a range of length scales, arises in the presence of continuous turnover since the inter-cluster regions can reassemble and repeatedly reach the percolation threshold, connecting neighboring clusters that will subsequently merge with each other. This stands in contrast with actomyosin networks with limited turnover, where once the network ruptures the local clusters are unable to reconnect ^16^.

Experimentally, we are able to induce local contraction both by increasing myosin activity (hence, increasing the contraction rate) or limiting filament length (and hence diminishing network connectivity) (Fig. 4). This is done using either Calyculin A, which enhances myosin activity by inhibiting myosin-light-chain phosphatase from dephosphorylating myosin, or by adding Capping Protein, which caps the barbed ends of free growing actin filaments. The addition of Calyculin, generates an intermediate state where within the same region we see alternating behaviors, with local contraction to a nearby contraction center, which subsequently merges with other clusters generating overall a global contraction pattern (Fig. 4a, Supplementary Video 11). The contraction pattern becomes more irregular at higher Calyculin concentrations. Similar behavior, with repeated transitions between local and global contractile behavior, is seen upon addition of intermediate levels of Capping Protein, which reduces network connectivity (Fig. 4b, left; Supplementary Video 12). At even higher Capping Protein concentration, the system breaks into local clusters (Fig. 4b, right). While these clusters can dynamically interact with each other and sometimes even merge, under these conditions, the contraction remains local and we do not observe significant coarsening or global contraction to a central cluster.

**Figure 4.**
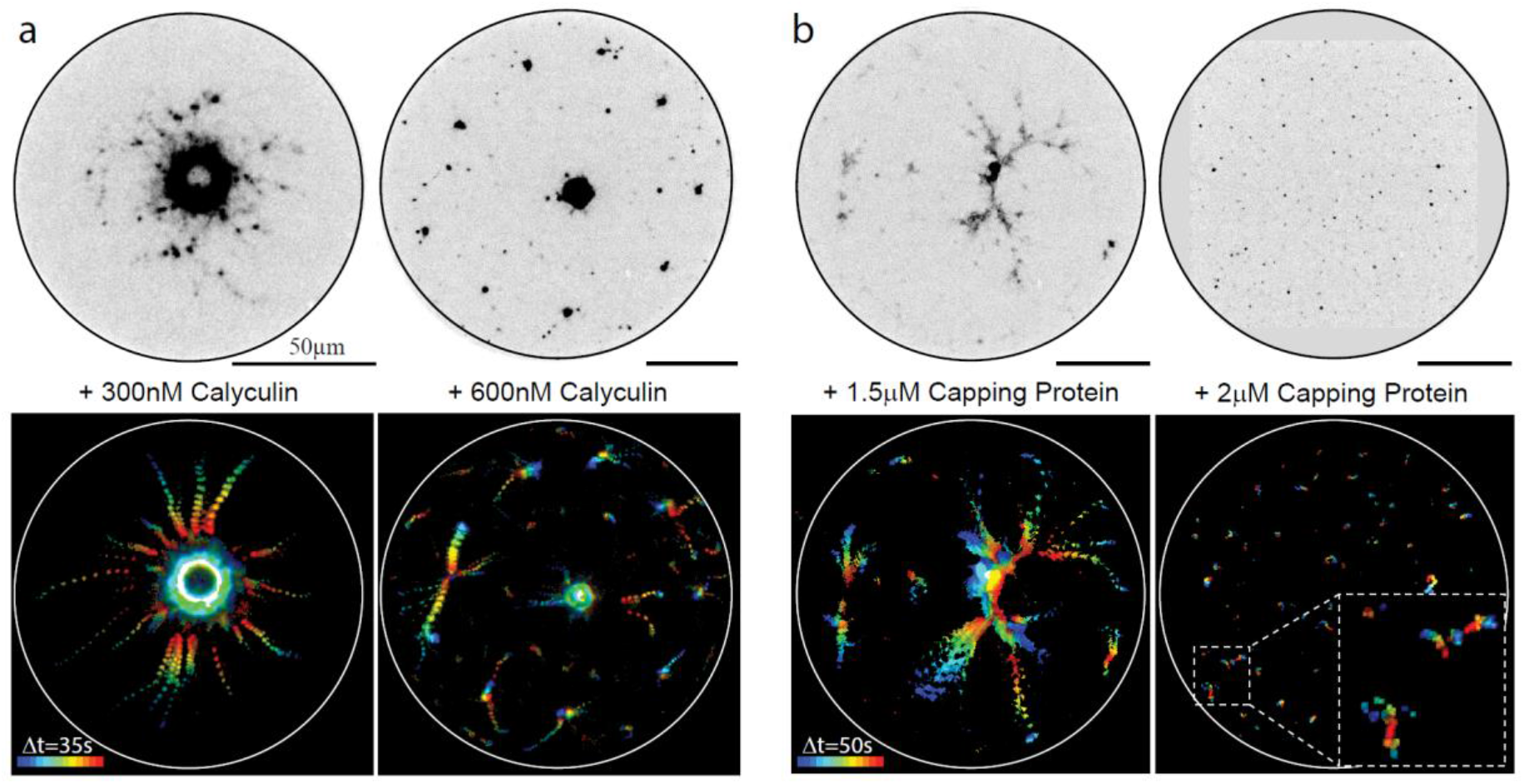
Transition from global to local contraction. Spinning disk confocal images of water-in-oil droplets containing 80% extract supplemented with (a) Calyculin A (Supplementary Video 11) or (b) Capping Protein (Supplementary Video 12). The dynamic contractile behavior is displayed by overlaying consecutive frames color-coded for time (bottom). The addition of Calylculin A or Capping protein leads to the formation of local contraction clusters. In some case, the local clusters eventually contract to a global contraction center, except at high enough Capping Protein concentration (b, right)

## DISCUSSION

Contracting actin networks exhibit a myriad of dynamic behaviors *in vivo*, including large-scale persistent flows, pulsatile contraction and local contractions ^1-3^. Many of these behaviors have been reconstituted *in vitro*, with systems exhibiting global contraction in the form of continuous flows ^23, 24^ or periodic waves ^13, 15, 27-29^, as well as local contraction into clusters ^12, 16, 33^. Here we use a reconstituted system based on cell extracts that exhibits rapid, physiological actin turnover rates, to explore a wide range of conditions and geometries, and reveal a size-dependent transition from continuous to periodic contraction.

The appearance of global contraction with different spatio-temporal patterns highlights the subtle interplay between network structure, contractile force generation and geometry. As discussed previously ^18, 34^, to generate and transmit forces across the system, the network must be percolated, yet these internal forces, in turn, also modulate network structure and connectivity. In the presence of rapid turnover, the extent of connected clusters within which forces are efficiently transmitted becomes a dynamic variable. Local clusters can form and contract but they can also join into an interconnected network, as actin assembly can form bridges between previously unconnected clusters. In this context, one has to consider a more dynamic version of the percolation problem, that considers internally generated large-scale advection as well as network assembly and disassembly. The contractile behavior of the system can no longer be summarized by a state diagram depending on the overall motor activity and network connectivity ^12, 13, 18, 35, 36^. Rather, one has to consider the geometry of the system and the spatial inhomogeneities that develop within the system. Our results suggest that the previously characterized state diagram is still relevant for describing the local contractile behavior at the mesoscale. However, to account for the emergent global behavior of the system, one has to consider the dynamic evolution of the system into regions with distinct mechanical properties. Interestingly, our work shows that the system size becomes an important factor in regulating the self-organized patterns that emerge; in particular, our work reveals a previously unappreciated transition whereby the same molecular components can self-organize into a persistent flow pattern or exhibit periodic waves, depending on the overall size of the system.

Recent work shows that similar contraction patterns containing distinct regions with different rheological properties emerge in cortical actomyosin networks of living cells, and provide the basis for a novel motility mechanism in disordered 3D environments ^37^. The unconnected gas-like state allows for the formation of protrusions at the leading edge of a cell that can penetrate into a disordered environment. Subsequent actin assembly generates a rigid, percolated cortical network which is pulled rearwards by myosin-generated contractile forces at cell rear. The suggested motility mechanism uses these rigid cortical regions, that assemble within the irregularly-shaped protrusions, to generate normal forces on the cells’ environment as the network is pulled rearwards, providing an efficient mechanism for momentum transfer to propel the cell forward.

The observed phenomenology in the cortex of these motile cells ^37^, with large-scale self-organized contractile patterns arising from a percolation transition in an active network undergoing continuous advection and turnover, is strikingly similar to our observations in bulk networks *in vitro*. In both cases the interconnected network formed above the rigidity-percolation transition facilitates long-range force transmission across the system (which is essential for global contraction) and at the same time is modulated by the internal advection generated by these forces. However, the geometry of the system has an important influence on the properties of the network. In the effective 1D geometry of the cortical networks on the surface of a cylindrical cell or bleb, the network can flow rearward as a solid, experiencing negligible stress, before fluidizing at the rear due to myosin-generated forces. However, in a 2D or 3D radially symmetric contracting network, the converging flow necessarily generates stress and hence the percolated network must be primarily viscous rather than solid (see more details in the SI).

The appearance of pulsatile modulation in actomyosin contraction is a common theme seen across many cell types in different contexts (e.g. ^6, 8, 10, 38^). While in some cases, the function of the pulsatile dynamics is unclear, in other cases the pulsatile dynamics are essential, enabling large-scale force generation while maintaining tissue integrity ^8, 9^, and facilitating cellular functions such as active transport and motility ^6^. Typically, these periodic or aperiodic waves emerge from complex nonlinear reactions in coupled mechano-chemical systems, involving regulators of the actomyosin machinery such as Rho GTPases ^10, 11, 36, 39, 40^. Our work shows that waves can also emerge in essentially mechanical systems, with turnover but without complex biochemical regulation. In this case, the periodic dynamics reflect an emerging spatiotemporal pattern that depends on the interplay between network contraction and turnover, rather than biochemical modulation of the actomyosin machinery. The period of the waves is comparable to actin turnover time scale, allowing the network in the wake of a wave front to reassemble and generate subsequent contracting wave fronts (see SI). Whether such mechanically-generated waves naturally arise in living systems and are functionally relevant remains to be seen.

Theoretically the phenomena of actomyosin contraction has been studied extensively using microscopic agent-based simulation as well as continuum hydrodynamic approaches. Several studies explicitly simulated microscopic models of actin filaments, myosin motors and crosslinkers (both motors and linkers stochastically bind to and unbind from filaments). Even without turnover, pulsed geometrically irregular contraction can emerge because of the positive feedback between local network density and effective contractile stress (due to more motors binding to denser actin arrays) ^19, 35^. Note that repeated irregular contractions in such systems depend on stochastic unbinding of motors and crosslinkers, which effectively dissolves transient network aggregates allowing for contraction to start elsewhere. Large-scale geometrically irregular pulsed contractions of similar nature appear in models with actin turnover ^21, 41^. Interestingly, a couple of models showed that such pulsed contractions can, in fact, be suppressed by actin turnover ^20, 42^.

Other models, albeit not microscopic, demonstrated that not just irregular, but also periodic and wavy contractions can develop in actomyosin networks. First, such periodic contractions can be generated by an effective relaxation oscillator based on coupling of network turnover with a highly nonlinear stress-density relation characterized by both a contractile phase at low network density and a relatively exotic swelling phase at high density ^30, 31^. Second, traveling contraction waves appear in dense actomyosin structures with myosin strain-dependent binding/unbinding kinetics ^43^, so in effect, some additional nonlinear kinetics has to be added to facilitate the existence of such waves. Third, one can build a spatial-temporal oscillator by coupling not just viscous but also elastic elements to contractile actomyosin network with turnover ^44^, which could be more relevant to supracellular, rather than cellular systems. Last, but not least, spatial-temporal instabilities in highly contractile actomyosin network with turnover, like those predicted by our model for large contraction rates (see the SI) were predicted in ^45^. None of these models however addressed the relationship between the contraction pattern and the geometry of the system, as done in this work.

Interestingly, our system also exhibits contraction in the form of spiral wave patterns (Fig. 1d, S2). Dynamic spiral patterns were observed in several physiological systems including the actin cortex, and are usually associated with highly nonlinear excitable systems ^38^ or intrinsic chirality at the molecular scale which is propagated to cellular scales ^46, 47^. While we have not yet attempted to quantitatively model spiral formation (our current model is limited to radially symmetric 2D patterns), we hypothesize that the spiral contraction waves we observe originate from a different and somewhat simpler mechanism. Along each radial direction, we have shown that the periodic waves are triggered, effectively, by an ‘integrate-and-fire’ oscillator ^48^: at the beginning of each cycle, the network density assembles to the percolation/contraction threshold (‘integrate’ part), which triggers a contraction that wipes out the density from the periphery (‘fire’ part) and resets the cycle. Each oscillator is characterized by a phase – the timing of the start of the cycle. There are lateral mechanical connections in the network between neighboring radial directions, so effectively we can consider them as coupled oscillators. Mathematical models demonstrated that such coupling leads, most frequently, to two possible patterns ^48^: 1) synchronization of the oscillator phases, which geometrically corresponds to periodic waves in the form of concentric contracting circles, which is most often the case, or 2) incremental phase shift between nearest-neighbor oscillators, which geometrically corresponds to a spiral pattern arising from some initial conditions. Note that such models predict that the spiral will develop so that along each radial direction the new wave starts at the periphery exactly when the previous wave crest merges with the dense network near the center, which seems to be the case in our system (Figs. 1d,e, S2, S3). Further research will show if this coupled oscillator mechanism is behind the spiral contraction waves and if these waves have a physiological significance.

Living systems exhibit patterns at different length scales depending on the mechanism driving their organization. The most prevalent mechanisms are diffusion-based biochemical pattern formation, such as Turing patterns, and various mechano-chemical patterns ^49^. Here, we demonstrate the existence of an emergent length scale based on coupling between mechanics, turnover and geometry. This characteristic length scale defines the transition length above which the contractile behavior of the system changes in a qualitative manner and periodic behavior develops. The size-dependent contractile behavior that emerges from the interplay between network advection, percolation and force generation was not appreciated before, and highlights the importance of the system’s geometry on its self-organized dynamics: the geometry of the system is not only reflected in the shape of the contracting wave front, but also in its temporal dynamics. More generally, this work provides an example of the profound impact that geometry and boundary conditions can have on pattern formation processes in cells and tissues.

## Acknowledgements

We thank Liora Garion for help with extract preparation. We thank Matthieu Piel and Raphael Voituriez for stimulating discussions and sharing unpublished results. We thank Jean-Francois Joanny for helpful advice on the modeling. We thank Peter Lenart, Guy Bunin, Alex Tayar, Anna Frishman and Sunghan Ro for comments on the manuscript.

This work was supported by a grant from the United States-Israel Binational Science Foundation to KK and Bruce Goode (grant No. 2017158). MS and AM are supported by National Science Foundation grants DMS 1953430 and DMS 2052515.

## Material and Methods

### Cell extracts, proteins and reagents

Concentrated M-phase extracts were prepared from freshly laid Xenopus laevis eggs as previously described ^25, 50, 51^. Briefly, Xenopus frogs were injected with hormones to induce ovulation and laying of unfertilized eggs for extract preparation. The eggs from different frogs were pooled together and washed with 1× MMR (100mM NaCl, 2mM KCl, 1mM MgCl_2_, 2mM CaCl_2_, 0.1mM EDTA, 5mM Na-Hepes, pH7.8), at 16°C. The jelly envelope surrounding the eggs was dissolved using 2% cysteine solution (in 100mM KCl, 2mM MgCl_2_ and 0.1mM CaCl_2_, pH7.8). Finally, the eggs were washed with CSF-XB (10mM K-Hepes pH7.7, 100mM KCl, 1mM MgCl_2_, 5mM EGTA, 0.1mM CaCl_2_ and 50mM sucrose) containing protease inhibitors (10 μg/ml each of leupeptin, pepstatin and chymostatin). The eggs were then packed using a clinical centrifuge and crushed by centrifugation at 15,000g for 15min at 4°C. The crude extract (the middle yellowish layer out of three layers) was collected, supplemented with 50mM sucrose containing protease inhibitors (10 μg/ml each of leupeptin, pepstatin and chymostatin), snap-frozen as 10 μl aliquots in liquid N_2_ and stored at −80°C. Typically, for each extract batch, a few hundred aliquots were made. Different extract batches exhibit similar behavior qualitatively, but the contraction rate and transition length exhibit some variation. All comparative analysis between conditions was done using the same batch of extract, but similar trends were observed in all extracts examined.

ActA–His was purified from strain JAT084 of Listeria monocytogenes (a gift from J. Theriot, Stanford University) expressing a truncated actA gene encoding amino acids 1–613 with a COOH-terminal six-histidine tag replacing the transmembrane domain, as described in ^25, 51^. Actin filaments Capping Protein Capz containing two subunits, α-1 and β-1, was purified to a final concertation of 66μM in 10mM Tris8, 50mM NaCl, 0.5mM DTT and 20% Sucrose. GST-mDia was purified to a final concertation of 45μM in 10mM Tris7.5, 50mM KCl, 1mM EGTA, 1mM MgCl_2_, 1mM DTT, 20mM Glutathione and 15% Sucrose. Purified proteins were aliquoted, snap-frozen in liquid N_2_ and stored at −80 °C until use. Calylculin A (Sigma) was added at 0-600nM to the extract mix.

Actin networks were labelled with GFP–Lifeact (the construct was a gift from C. Field, Harvard Medical School). GFP–Lifeact was purified and concentrated to a final concentration of 252 μM in 100mM KCl, 1mM MgCl_2_, 0.1mM CaCl_2_, 1mM DTT and 10% sucrose and stored at −80 °C until use.

### Sample preparation

For samples with 80% extract, an aqueous mix was prepared by mixing the following: 8μl crude extract, 0.5 μl 20× ATP regenerating mix (150mM creatine phosphate, 20mM ATP, 20mM MgCl_2_), 0.5 μl of 10 μM GFP–Lifeact and any additional proteins as indicated. The final volume in was adjusted to 10 μl by adding XB buffer (10mM Hepes, 5mM EGTA, 100mM KCl, 2mM MgCl_2_ and 0.1mM CaCl_2_ at pH7.8). For samples with higher extract concentration (95-98%), crude extract was supplemented with a 50x concentrated mix containing ATP regenerating system and GFP–Lifeact to generate the same final concentrations.

Emulsions were made by adding 3% (v/v) extract mix to degassed mineral oil (Sigma) containing 4% cetyl PEG/PPG-10/1 dimethicone (Abil EM90, Evnok Industries) and stirring for 1min at 4°C. The mix was then incubated for an additional 10min on ice to allow the emulsions to settle.

Samples were made in chambers assembled from two passivated coverslips separated by 10μm or 30 μm-thick double stick tape (3M), sealed with wax (vaseline:lanolin:paraffin at 1:1:1 ratio) and attached to a glass slide. Passivation was done by incubating cleaned coverslips in silanization solution (5% dichlorodimethylsilane in heptane) for 20 minutes, washing in heptane, sonicating twice in DDW for 5 minutes and once in ethanol for 5 minutes. For capillary experiments, the extract mix (without making an emulsion) was introduced into a 50 × 500 μm rectangular capillary (CM Scientific) through capillary action and the capillary ends were sealed with VALAP. Samples were typically imaged 10-60 min after sample preparation.

### Microscopy

Emulsions were imaged on Zeiss Observer Z1 confocal microscope fitted with a Yokogawa CSU-X1 spinning-disk using Slidebook software for acquisition at room temperature (Intelligent Imaging Innovations). Samples were illuminated with 488nm laser and images were captured at the mid-plane of the sample, so the network velocity is mostly within the imaging plane.

High temporal resolution images to characterize network properties were obtained with 10x air (NA = 0.5), 20x air (NA = 0.75) or 40x oil (NA = 1.3) objectives with or without optovar (1.6x), depending on the size of the droplets. Images were acquired onto a 512×512 EM-CCD camera (QuantEM; Photometrix) at 0.5s time resolution.

Low time resolution images for characterizing the global contractile behavior in a population of droplets were acquired using a 2048×2048 CMOS camera (Zyla, Andor) with a 20x air (NA = 0.5) objective. Images were acquired with 2×2 binning at 15s time interval for a duration of 8min.

### Analysis

#### Network properties of bulk actin network in emulsions

High time resolution time-lapse videos were acquired at the mid-plane of the sample to obtain the bulk actin network flows as in ^23^. Time-lapse videos were background-corrected, corrected for uneven illumination by normalizing with flat field corrections and bleach corrected using an exponential fit to the total image intensity as a function of time.

The velocity fields were extracted from the corrected time-lapse videos using either optical flow with a reaction term to account for network turnover using the code adapted from ^52^ and modified as described below, or particle image velocimetry (PIV) as described previously ^23^. Optical flow analysis was done on movies with 0.5s time interval, whereas PIV analysis was done at 2.5-10s time interval with additional averaging of the spatio-temporal correlation function over 3-40 consecutive frames ^23^. For continuous contraction, both methods yielded similar results. However, for the non-steady contraction, only the optical flow method could be used because the temporal changes in the velocity field were fast compared to the time scales used for averaging the spatio-temporal correlation function in the PIV method (which are essential for obtaining a reasonable velocity field).

Optical flow analysis was performed on 15×15 pixel window. Consecutive pairs of images were first smoothed with a Gaussian filter (sigma 1.5 pixels), and subsequently analyzed with a reaction term using the Matlab code provided in ^52^. The velocity fields were subsequently averaged over 5 consecutive frames (=2.5s) using a 15×15 median filter to reduce noise. For continuous contraction, the velocity field was further down sampled to a final resolution of 12.5s and 15×15 pixels by averaging. The contraction rate in the continuous contraction regime was determined from the slope of the linear fit to the radial velocity as a function of distance to the contraction center ^23^.

Radial kymographs of the density and the velocity fields as a function of distance from the contraction center were obtained by angular averaging. The averaging was done over all angles for droplets with a symmetric contraction or over a smaller manually selected angular range otherwise. Angular kymographs (for spiral movies) were determined by considering the angular variation in a specified radial range.

#### Population statistics

The population statistics were acquired from time-lapse images of droplets with varying sizes at 15s time interval (to capture at least 2 frames per period in the periodic regime). For each droplet, radial intensity kymographs were obtained as above. The state of contraction in each droplet was categorized as continuous or periodic based on manual inspection of the radial kymographs. For droplets exhibiting periodic density modulations (i.e. parallel diagonal lines in the radial kymograph), the time period was determined from analyzing the time dependent intensity variation for a small radial region (i.e. along a vertical line in the kymograph) using in-bulit Matlab function ‘periodogram’, where the periodicity is determined from the position of the most prominent peak in the Fourier spectrum, estimated from the location of the center of a best-fit Gaussian. The width of the best-fit Gaussian is used to estimate the error in the periodicity. The period was determined only for droplets that remained stationary throughout the movie. Droplets which exhibit irregular density modulations rather than a clear periodic pattern were characterized to be in a mixed state.

The transition length for various conditions was determined from considering the contraction state vs size (Fig. 2b). For conditions where we observe a size range in which droplets were found exhibiting both continuous and periodic contraction, the transition length (*R*_*tr*_) was defined to minimize the number of droplets exhibiting periodic contraction with R<*R*_*tr*_ and continuous contraction with R>*R*_*tr*_. If there was no overlap, the transition length is reported as mean of the sizes of the smallest droplet exhibiting periodic contraction and the largest continuous exhibiting continuous contraction droplet.

The spectral time-lapse (STL) images for different concentrations of added Capping Protein and Calyculin (Fig. 4) were obtained using the STL algorithm ^53^. The time-lapse images were masked with either a moments binary mask from Fiji or a manual intensity threshold and time was color-coded using STL algorithm over 35 – 50s as indicated, depending on the rate of network displacements.

### Modeling

To model the actomyosin network, we use a system of transport (assembly, disassembly and drift) and force-balance (balancing myosin-generated contractile and crosslinked-actin viscous stresses) equations. These two partial differential equations, with boundary conditions specified in the SI, determine the network’s density and drift velocity, which are both functions of space and time. In the continuous model, the effective contractile stress and viscosity are functions of the network density as specified in the SI. We solve the model numerically, with parameters obtained from the data, in 1D and in a radially symmetric 2D case. We also simulate numerically a 1D stochastic agent-based model in which material nodes of the network appear and disappear at random times and locations with constant rates. The neighboring node pairs do not interact if the distance between them is greater than the percolation length, stay at constant distance from each other if this distance is less than the percolation length but greater than the contraction length, and converge with a constant rate if their mutual distance is less than the contraction length. Further details are provided in the SI.

## References

1. Field, C.M. & Lenart, P. Bulk cytoplasmic actin and its functions in meiosis and mitosis. Current Biology 21, R825–R830 (2011).

2. Salbreux, G., Charras, G. & Paluch, E. Actin cortex mechanics and cellular morphogenesis. Trends in Cell Biology 22, 536–545 (2012).

3. Munjal, A. & Lecuit, T. Actomyosin networks and tissue morphogenesis. Development 141, 1789–1793 (2014).

4. Koenderink, G.H. & Paluch, E.K. Architecture shapes contractility in actomyosin networks. Current Opinion in Cell Biology 50, 79–85 (2018).

5. Murrell, M., Oakes, P.W., Lenz, M. & Gardel, M.L. Forcing cells into shape: the mechanics of actomyosin contractility. Nature Reviews Molecular Cell Biology 16, 486–498 (2015).

6. Munro, E., Nance, J. & Priess, J.R. Cortical flows powered by asymmetrical contraction transport PAR proteins to establish and maintain anterior-posterior polarity in the early C. elegans embryo. Developmental Cell 7, 413–424 (2004).

7. Lenart, P. et al. A contractile nuclear actin network drives chromosome congression in oocytes. Nature 436, 812–818 (2005).

8. Martin, A.C., Kaschube, M. & Wieschaus, E.F. Pulsed contractions of an actin-myosin network drive apical constriction. Nature 457, 495–499 (2009).

9. Jodoin, J.N. et al. Stable force balance between epithelial cells arises from F-actin turnover. Developmental Cell 35, 685–697 (2015).

10. Nishikawa, M., Naganathan, S.R., Julicher, F. & Grill, S.W. Controlling contractile instabilities in the actomyosin cortex. eLife 6, e19595 (2017).

11. Agarwal, P. & Zaidel-Bar, R. Principles of actomyosin regulation in vivo. Trends in Cell Biology 29, 150–163 (2019).

12. Backouche, F., Haviv, L., Groswasser, D. & Bernheim-Groswasser, A. Active gels: dynamics of patterning and self-organization. Physical Biology 3, 264 (2006).

13. Bendix, P.M. et al. A quantitative analysis of contractility in active cytoskeletal protein networks. Biophysical Journal 94, 3126–3136 (2008).

14. Reymann, A.-C. et al. Actin network architecture can determine myosin motor activity. Science 336, 1310–1314 (2012).

15. Kohler, S. & Bausch, A.R. Contraction mechanisms in composite active actin networks. PLoS One 7, e39869 (2012).

16. Alvarado, J., Sheinman, M., Sharma, A., MacKintosh, F.C. & Koenderink, G.H. Molecular motors robustly drive active gels to a critically connected state. Nature Physics 9, 591–597 (2013).

17. Ennomani, H. et al. Architecture and connectivity govern actin network contractility. Current Biology 26, 616–626 (2016).

18. Alvarado, J., Sheinman, M., Sharma, A., MacKintosh, F.C. & Koenderink, G.H. Force percolation of contractile active gels. Soft Matter 13, 5624–5644 (2017).

19. Belmonte, J.M., Leptin, M. & Nedelec, F. A theory that predicts behaviors of disordered cytoskeletal networks. Molecular Systems Biology 13, 941 (2017).

20. McFadden, W.M., McCall, P.M., Gardel, M.L. & Munro, E.M. Filament turnover tunes both force generation and dissipation to control long-range flows in a model actomyosin cortex. PLoS Computational Biology 13, e1005811 (2017).

21. Hiraiwa, T. & Salbreux, G. Role of turnover in active stress generation in a filament network. Physical Review Letters 116, 188101 (2016).

22. Jansen, S., Johnston, A., Eskin, J. & Goode, B.L. Mechanisms controlling cellular actin disassembly and turnover. Nature Reviews of Molecular and Cell Biology (2018).

23. Malik-Garbi, M. et al. Scaling behaviour in steady-state contracting actomyosin networks. Nature Physics 15, 509–516 (2019).

24. Pinot, M. et al. Confinement induces actin flow in a meiotic cytoplasm. Proc Natl Acad Sci U S A 109, 11705–11710 (2012).

25. Abu-Shah, E. & Keren, K. Symmetry breaking in reconstituted actin cortices. eLife 3, e01433 (2014).

26. Tan, T.H. et al. Self-organization of stress patterns drives state transitions in actin cortices. Science Advances 4, eaar2847 (2018).

27. Ezzell, R.M., Brothers, A.J. & Cande, W.Z. Phosphorylation-dependent contraction of actomyosin gels from amphibian eggs. Nature 306, 620–622 (1983).

28. Field, C.M. et al. Actin behavior in bulk cytoplasm is cell cycle regulated in early vertebrate embryos. J Cell Sci 124, 2086–2095 (2011).

29. Sakamoto, R. et al. Tug-of-war between actomyosin-driven antagonistic forces determines the positioning symmetry in cell-sized confinement. Nature Communications 11, 1–13 (2020).

30. Pohl, T. Periodic contraction waves in cytoplasmic extracts, in Biological Motion, Vol. 89 85–94 (Springer, 1990).

31. Alt, W. & Dembo, M. Cytoplasm dynamics and cell motion: two-phase flow models. Mathematical biosciences 156, 207–228 (1999).

32. Ierushalmi, N. et al. Centering and symmetry breaking in confined contracting actomyosin networks. eLife 9, e55368 (2020).

33. Wollrab, V. et al. Polarity sorting drives remodeling of actin-myosin networks. Journal of Cell Science 132, jcs219717 (2019).

34. Bueno, C., Liman, J., Schafer, N.P., Cheung, M.S. & Wolynes, P.G. A generalized Flory-Stockmayer kinetic theory of connectivity percolation and rigidity percolation of cytoskeletal networks. PLoS Computational Biology 18, e1010105 (2022).

35. Freedman, S.L., Hocky, G.M., Banerjee, S. & Dinner, A.R. Nonequilibrium phase diagrams for actomyosin networks. Soft Matter 14, 7740–7747 (2018).

36. Banerjee, S., Gardel, M.L. & Schwarz, U.S. The actin cytoskeleton as an active adaptive material. Annual Review of Condensed Matter Physics 11, 421–439 (2020).

37. García-Arcos, J.M. et al. Advected percolation in the actomyosin cortex drives amoeboid cell motility. bioArxiv (2022).

38. Bement, W.M. et al. Activator-inhibitor coupling between Rho signalling and actin assembly makes the cell cortex an excitable medium. Nature Cell Biology 17, 1471–1483 (2015).

39. Allard, J. & Mogilner, A. Traveling waves in actin dynamics and cell motility. Current Opinion in Cell Biology 25, 107–115 (2013).

40. Staddon, M.F., Munro, E.M. & Banerjee, S. Pulsatile contractions and pattern formation in excitable actomyosin cortex. PLoS Computational Biology 18, e1009981 (2022).

41. Mak, M., Zaman, M.H., Kamm, R.D. & Kim, T. Interplay of active processes modulates tension and drives phase transition in self-renewing, motor-driven cytoskeletal networks. Nature Communications 7, 1–12 (2016).

42. Yu, Q., Li, J., Murrell, M.P. & Kim, T. Balance between force generation and relaxation leads to pulsed contraction of actomyosin networks. Biophysical journal 115, 2003–2013 (2018).

43. Banerjee, D.S., Munjal, A., Lecuit, T. & Rao, M. Actomyosin pulsation and flows in an active elastomer with turnover and network remodeling. Nature Communications 8, 1121 (2017).

44. Dierkes, K., Sumi, A., Solon, J.e.o. & Salbreux, G. Spontaneous oscillations of elastic contractile materials with turnover. Physical Review Letters 113, 148102 (2014).

45. Hannezo, E., Dong, B., Recho, P., Joanny, J.-F.c. & Hayashi, S. Cortical instability drives periodic supracellular actin pattern formation in epithelial tubes. PNAS 112, 8620–8625 (2015).

46. Naganathan, S.R., Furthauer, S., Nishikawa, M., Julicher, F. & Grill, S.W. Active torque generation by the actomyosin cell cortex drives left-right symmetry breaking. eLife 3, e04165 (2014).

47. Tee, Y.H. et al. Cellular chirality arising from the self-organization of the actin cytoskeleton. Nature Cell Biology 17, 445–457 (2015).

48. Bressloff, P.C. Mean-field theory of globally coupled integrate-and-fire neural oscillators with dynamic synapses. Physical Review E 60, 2160 (1999).

49. Bailles, A.i., Gehrels, E.W. & Lecuit, T. Mechanochemical Principles of Spatial and Temporal Patterns in Cells and Tissues. Annual review of cell and developmental biology 38 (2022).

50. Field, C.M., Nguyen, P.A., Ishihara, K., Groen, A.C. & Mitchison, T.J. Xenopus egg cytoplasm with intact actin, in Methods in enzymology, Vol. 540 399–415 (Elsevier, 2014).

51. Abu-Shah, E., Malik-Garbi, M. & Keren, K. Reconstitution of actin cortices, in Building a cell from its component parts. (eds. J. Ross & W. Marshall) (Elsevier, 2014).

52. Vig, D.K., Hamby, A.E. & Wolgemuth, C.W. On the quantification of cellular velocity fields. Biophys J 110, 1469–1475 (2016).

53. Madan, C.R. & Spetch, M.L. Visualizing and quantifying movement from pre-recorded videos: the spectral time-lapse (STL) algorithm. F1000Research 3 (2014).

