## Supplementary Information for "Size-dependent transition from steady contraction to waves in actomyosin networks with turnover"

### Supplementary Text

#### *Continuous model*

To model the actomyosin network, we turn to the basic transport (1) and force-balance (2) equations that have been used previously [1-4]:

$$\frac{\partial\rho}{\partial t} = \alpha - \beta\rho - \nabla\cdot(\vec{u}\rho) \quad (1)$$

$$\nabla\cdot\sigma = 0 \quad (2)$$

Here $\rho$ is the network density, $\alpha$ is the assembly rate of the network, $\beta$ is the disassembly rate, and $\vec{u}$ is the network velocity. The first equation is the mass conservation law for the drifting and turning over network. The velocity here is a dynamic variable that is a function of the spatial coordinate and time, and must be found using Eq. 2. Also, the assembly and disassembly rates need not be constant and can be functions of the density.

In the second equation, $\sigma$ is the total network internal stress tensor. Usually, the force balance equation is written in the form: $\nabla\cdot\sigma = \zeta\vec{u}$ , where $\zeta$ is the effective drag between the network and the fluid in which the network is immersed. However, we demonstrated earlier [2,3] that the friction between the network and fluid is much smaller than the effective contractile and internal viscous forces within the network, so we use the approximate Eq. 2 instead. For simplicity and in the absence of rheological measurements, we assume that the network is a very viscous isotropic Newtonian fluid [5], albeit compressible (which is obvious from the variable network density). To be clear, the combination of the network and solute that it is immersed in is incompressible, but by neglecting the solute fraction we may consider the network as a simple compressible fluid.

Thus, the components of the total stress tensor are:

$$\sigma_{ij} = \underbrace{\mu\left(\frac{\partial u_i}{\partial x_j} + \frac{\partial u_j}{\partial x_i}\right) - \left(\frac{2}{3}\mu - \lambda\right)(\nabla\cdot\vec{u})\delta_{ij}}_{viscous} + \underbrace{\sigma_{contr}\delta_{ij}}_{contractile} \quad (3)$$

Here, the last term represents the active myosin-powered contraction, where the scalar $\sigma_{contr}$ is the magnitude of the contractile stress. The other terms describe the passive stress of the actin network deformations. The parameters $\mu$ and $\lambda$ are the effective shear and bulk viscosities, respectively. Taking the divergence of the total stress provides us with the force-balance equation:

$$\mu\nabla^2\vec{u} + \left(\frac{1}{3}\mu + \lambda\right)\nabla(\nabla\cdot\vec{u}) + \nabla\sigma_{contr} = 0 \quad (4)$$

The density dependencies of the viscosity and the contractile stress are crucial for the large-scale network dynamics. A central feature of our model is the consideration of the coexistence of distinct mechanical regimes characterized by a different viscosity and active stress in different regions of the system. We adopt the state diagram from [6] in the following interpretation to describe the local state in each region (Fig. 3a). The state of the network in [6] is determined by two factors – connectivity and contractility. We make the following simplifications: 1) the connectivity is proportional to the local actin network density, $\rho$ ; 2) the contractile stress is an increasing function of the local myosin density; 3) the myosin density is an increasing function of the actin network density; 4) myosin kinetics is fast compared to actin turnover, meaning that the local myosin density adjusts rapidly to the local actin density. This fourth assumption allows us to describe the system by considering a single (actin) density field. The behavior of the system is then restricted to a straight line (or a curve) on the state diagram (Fig. 3a) – passing through the unconnected (I), interconnected (II) and contractile (III) states, as this single density varies. This assumption is not necessarily true in our experimental system – myosin kinetics may not be fast – but qualitatively this would not change the system's behavior. The main point is that locally the system is in one of these three distinct mechanical states. Making the simplification of the fast myosin kinetics allows to consider only one density field explicitly (instead of two), and describe the local state of the system as a function of the local density.

We assume that the network in the unconnected state has zero viscosity and contractility, in the interconnected/non-contractile, state – non-zero viscosities and zero contractility, and in the interconnected/contractile state – non-zero viscosities and contractility. More specifically,

$$\mu = \begin{cases} 0 & \rho < \rho_{12} \\ \mu_0 & \rho > \rho_{12} \end{cases}, \lambda = \begin{cases} 0 & \rho < \rho_{12} \\ \lambda_0 & \rho > \rho_{12} \end{cases}, \sigma_{contr} = \begin{cases} 0 & \rho < \rho_{23} \\ \sigma_0 & \rho > \rho_{23} \end{cases} \tag{5}$$

where $\rho_{12}$ and $\rho_{23}$ are the density thresholds for interconnectedness and contractility, respectively (Fig. 3a) and $\rho_{12} < \rho_{23}$. Note that we assume that above the interconnected/contractile threshold, the viscosities and contractility are constants. This assumption is not crucial, but greatly simplifies the model. This density independence also leads to the telescopic character of the contraction [2]: the inward velocity increases almost linearly as a function of the distance from the contraction center. Note also that from now on, the force balance equations are applicable only to the interconnected network, for $\rho > \rho_{12}$. Wherever $\rho < \rho_{12}$, the velocity is equal to zero, and the transport equation becomes an ordinary differential equation as a function of time.

We discuss the boundary conditions for the model in detail below; here, we mention briefly that the boundary conditions for the force balance equation (Eq. 4) are: 1) zero velocity at the inner boundary of the interconnected network (at the boundary of the aggregate that forms at the contraction center [3]); 2) zero normal component of the total stress (Eq. 3) at the outer boundary of the interconnected network.

We consider two geometries: 1) one-dimensional (1D) or, 2) radially symmetric two-dimensional (2D). In the 1D case, the force balance equation after integration simply states that the total internal network stress tensor is equal to a constant. Because of the boundary condition of zero stress at the outer boundary of the network, this constant is equal to zero. Thus, in 1D, Eqs. 1 and 4 reduce to:

$$\frac{\partial \rho}{\partial t} = \alpha - \beta\rho - \frac{\partial}{\partial x}(u\rho) \tag{6}$$

$$\eta_0 \frac{\partial u}{\partial x} + \sigma_{contr} = 0 . \qquad (7)$$

Here $\eta_0 = \frac{4}{3}\mu_0 + \lambda_0$ is the effective viscosity coefficient in the 1D model, $x$ is the 1D coordinate, $u$ is the 1D network velocity.

For $\rho > \rho_{23}$, Eq. 7 becomes: $\frac{\partial u}{\partial x} = -k \to u = u_0 - kx$, where $u_0$ is a constant and $k = \frac{\sigma_0}{\eta_0}$. The parameter $k$ is the observable contraction rate. In the unconnected phase, the velocity is simply equal to zero. In the interconnected/non-contractile state, $\rho_{12} < \rho < \rho_{23}$, where the viscosity is non-zero but the contractility is zero, Eq. 7 becomes: $\frac{\partial u}{\partial x} = 0 \to u = u_0$. To summarize, the 1D model becomes:

$$\frac{\partial \rho}{\partial t} = \alpha - \beta\rho - \frac{\partial}{\partial x}(u\rho), u = \begin{cases} 0 & \rho < \rho_{12} \\ const & \rho_{12} < \rho < \rho_{23} \\ const - kx & \rho > \rho_{23} \end{cases} \qquad (8)$$

The constants are found from the boundary condition near the center of the droplet, $u(0) = 0$, and the condition of the continuity of the velocity.

For example, when $\rho > \rho_{23}$ at $0 < x < r_{23}$, $\rho_{12} < \rho < \rho_{23}$ at $r_{23} < x < r_{12}$, and $\rho_{12} > \rho$ at $r_{12} < x < R$ (where $r_{23}$ and $r_{12}$ are defined as where $\rho(r_{23}) = \rho_{23}$ and $\rho(r_{12}) = \rho_{12}$, respectively), then $u = -kx$ at $0 < x < r_{23}$, $u = -kr_{23}$ at $r_{23} < x < r_{12}$, and $u = 0$ at $r_{12} < x < R$. In principle, there could be multiple regions of interspersed contractile and interconnected/non-contractile network (see the general formulas for the 2D case below). In these cases, the expansion of Eq. 8 is straightforward.

In 2D, for the radially symmetric case, the transport equation becomes:

$$\frac{\partial \rho}{\partial t} = \alpha - \beta\rho - \frac{1}{r}\frac{\partial}{\partial r}(ru\rho), \qquad (9)$$

while the force balance equation is:

$$\mu_0 \left( \frac{1}{r}\frac{\partial}{\partial r}\left( r\frac{\partial u}{\partial r} \right) - \frac{u}{r^2} \right) + \left( \frac{\mu_0}{3} + \lambda_0 \right) \frac{\partial}{\partial r}\left( \frac{1}{r}\frac{\partial (ru)}{\partial r} \right) + \frac{\partial \sigma_0}{\partial r} = 0 . \qquad (10)$$

Here $r$ is the radial coordinate in a polar coordinate system (centered at the center of the aggregate within the droplet), and $u(r,t)$ is the radial component of the velocity. The radial component of the stress tensor in this case has the form:

$$\sigma_{rr} = 2\mu_0 \frac{\partial u}{\partial r} - \left( \frac{2}{3}\mu_0 - \lambda_0 \right) \frac{1}{r}\frac{\partial}{\partial r}(ru) + \sigma_0 . \qquad (11)$$

In 2D, as the active stress is assumed to be piece-wise constant, $\sigma_{contr} = const$ if $\rho \neq \rho_{12}, \rho_{23}$, the force balance (Eq. 10) within each region becomes:

$\mu_0\left(\frac{1}{r}\frac{\partial}{\partial r}\left(r\frac{\partial u}{\partial r}\right)-\frac{u}{r^2}\right)+\left(\frac{\mu_0}{3}+\lambda_0\right)\frac{\partial}{\partial r}\left(\frac{1}{r}\frac{\partial(ru)}{\partial r}\right)=0$. Simple calculations allow to re-write the left-hand side in a particularly convenient form: $\eta_0\left(\frac{\partial^2 u}{\partial r^2}+\frac{1}{r}\frac{\partial u}{\partial r}-\frac{u}{r^2}\right)=0$, where $\eta_0=\frac{4}{3}\mu_0+\lambda_0$. The general solution of this equation has the form: $u=ar+b/r$, where $a$ and $b$ are arbitrary constants (for any given time moment) to be found from the boundary conditions. This general form of the velocity profile is applicable both in the contractile $\rho>\rho_{23}$ and in the interconnected/non-contractile $\rho_{12}<\rho<\rho_{23}$ states. In the unconnected phase the velocity is simply zero. Substituting this expression for the radial velocity into Eq. 11, we obtain the general formula for the radial stress:
$\sigma_{rr}=\left(\frac{2\mu_0}{3}+2\lambda_0\right)a-2\mu_0 b/r^2+\sigma_0$.
To summarize, the 2D model becomes:

$$\frac{\partial\rho}{\partial t}=\alpha-\beta\rho-\frac{1}{r}\frac{\partial}{\partial r}(ru\rho),\quad u=\begin{cases}0 & \rho<\rho_{12}\\ (const)r+const/r & \rho_{12}<\rho<\rho_{23}\\ (const)r+const/r & \rho>\rho_{23}\end{cases}\qquad (12)$$

Note that the constants in the expression for the velocities are different in each region (as described below, they are determined by stitching the results in the different regions while taking into account the boundary conditions).

Here are the remaining model assumptions and boundary conditions, after which the particular solution for the 2D velocity profile is outlined further:

1. A boundary condition for the velocity: $u(0)=0$ in 1D or $u(r_0)=0$ in 2D, meaning that at its innermost boundary the network sticks to the aggregate that forms at the contraction center [3].
2. A second boundary condition: no-stress at the outer interconnected network boundary $x=r_{12}$ in 1D or $r=r_{12}$ in 2D, where $r_{12}$ is the point at which the interconnected network ends (so that $\rho(r_{12})=\rho_{12}$). This is already applied in Eq. 8 for 1D. In 2D, this appears as zero radial stress at the outer interconnected network boundary: $\sigma_{rr}(r_{12})=0$.
3. A continuity condition for the velocity: $u(x)$ in 1D or $u(r)$ in 2D are continuous functions for $\rho>\rho_{12}$. Additionally, in 2D, a continuity condition for the radial stress: $\sigma_{rr}(r)$ is a continuous function for $\rho>\rho_{12}$.
4. At the outer boundary of the interconnected network ($\rho=\rho_{12}$), the network boundary grows at constant rate equal to $v_0$. Mathematically, the boundary moves with a velocity: $u(x)+v_0$ in 1D or $u(r)+v_0$ in 2D for $x$ and $r$ such that $\rho(x)=\rho_{12}$ or $\rho(r)=\rho_{12}$, respectively. Physically, the growth of the network boundary can be explained by the polymerization and elongation of filaments connected to the network at the edge of the network.
5. $\alpha$, the network assembly rate, is assumed to be density dependent, so that the rate saturates to a constant at larger densities and decreases to a smaller constant at lower densities. Such density dependent assembly could arise e.g., from the contribution of autocatalytic, Arp2/3-dependent, assembly. We use the expression: $\alpha=\alpha_0+(\bar{\alpha}-\alpha_0)(1-\exp(-\rho/\rho_0))$.

The density dependency of the assembly rate is not critical. Qualitatively, the model works well without this assumption, but the results are more numerically robust with the assumed density dependence.

6. We solve the equations in 1D on the domain $0 < x < R$, where R is the droplet radius, and in 2D on $r_0 < r < R$.

The particular solution for the velocity profile in 2D depends on the breakdown of the domain $r_0 < r < R$ into the different regimes: contractile, interconnected/non-contractile, and unconnected. We consider the solution in the part of the domain where $r_0 < r < r_{12}$, having defined $r_{12}$ as the first point from $r = r_0$ where $\rho \leq \rho_{12}$ (the first point where the network is no longer connected). In principle, there can be a situation in which two interconnected and/or contractile network regions are separated by an unconnected region. In that case, one has to analyze each interconnected region separately. In our simulations, we encountered cases in which an interconnected but not contractile network is separated by an unconnected region from another interconnected and contractile region. In these case, the situation is simple: the velocity in the interconnected/non-contractile region, separated by the unconnected region from the contractile network, is equal to zero. The analysis becomes nontrivial if there are multiple contractile networks separated by unconnected networks. In our simulations, we have not encountered such cases, and so we omit such analysis here.

We consider the case of one interconnected domain $r_0 < r < r_{12}$. We divide this domain into regions with distinct mechanical states. In region $i$ ($i = 1, 2, \ldots, n$), which includes the part of the domain $r^{i-1,i} \leq r < r^{i,i+1}$, the velocity profile is $u_i(r) = a_i r + b_i/r$. Note that when $i = 1$, $r^{0,1} = r_0$ and when $i = n$, $r^{n,n+1} = r_{12}$. Numerical solutions show that region $i = 1$ is always contractile, as the density is always greater than $\rho_{23}$ near the inner boundary of the domain.

The linear system for the coefficients $\{a_i, b_i\}_{i=1}^{n}$ (total $2n$-many) requires $2n$-many equations to be solved. Two come from the boundary conditions:

$$u_1(r_0) = a_1 r_0 + b_1/r_0 = 0 \quad (13)$$

(zero velocity at the inner boundary) and

$$\sigma_{rr}(r_{12}) = 0 = \begin{cases} \left(\dfrac{2\mu_0}{3} + 2\lambda_0\right) a_n - 2\mu_0 b_n / r_{12}^2, & n \text{ even} \\ \left(\dfrac{2\mu_0}{3} + 2\lambda_0\right) a_n - 2\mu_0 b_n / r_{12}^2 + \sigma_0, & n \text{ odd} \end{cases} \quad (14)$$

(no radial stress at the outer boundary). Note that when $n$ is even, the final region is non-contractile and when $n$ is odd, the final region is contractile.

The remaining equations come from the continuity conditions applied across each point $r^{i,i+1}$. For each $i = 1, 2, \ldots, n-1$, velocity continuity requires that

$$a_i r^{i,i+1} + b_i/r^{i,i+1} = a_{i+1} r^{i,i+1} + b_{i+1}/r^{i,i+1} \quad (15)$$

and continuity of $\sigma_{rr}$ requires that

$$\begin{cases} \left(\frac{1}{3}+\frac{\lambda_0}{\mu_0}\right)a_i - b_i/\left(r^{i,i+1}\right)^2 = \left(\frac{1}{3}+\frac{\lambda_0}{\mu_0}\right)a_{i+1} - b_{i+1}/\left(r^{i,i+1}\right)^2 + k, & i \text{ even} \\ \left(\frac{1}{3}+\frac{\lambda_0}{\mu_0}\right)a_i - b_i/\left(r^{i,i+1}\right)^2 + k = \left(\frac{1}{3}+\frac{\lambda_0}{\mu_0}\right)a_{i+1} - b_{i+1}/\left(r^{i,i+1}\right)^2, & i \text{ odd} \end{cases} \quad (16)$$

Here $k=\sigma_0/2\mu_0$ is the approximate contraction rate for the 2D model (which turns out to be the same as that in the 1D case; see below). These make up in total the $2n$ equations necessary to solve for the $2n$-many coefficients for the piece-wise continuous velocity profile. Numerical solutions demonstrate two interesting features of this system (see the simulation movies and snapshots from the simulations in Fig. 3): 1) in the interconnected/non-contractile regions, the velocity profile is relatively close to being flat. 2) in the contractile regions, the velocity profile is relatively close to being linear, with a slope equal to $k=\sigma_0/2\mu_0$ multiplied by two factors, both of which are on the order of unity. The first factor is a function of the two viscosities, $\lambda_0$ and $\mu_0$ (it is of order unity if $\lambda_0$ and $\mu_0$ are of the same order of magnitude). In what follows, we consider a specific case that gives the simplest result and choose $\lambda_0 = \frac{2}{3}\mu_0$. Then, the factor $\left(\frac{1}{3}+\frac{\lambda_0}{\mu_0}\right)$ in Eq. 16 and in the first factor for the contraction rate become equal to unity. In this specific case, also $\eta_0 = 2\mu_0$, and the contraction rates in 1D and 2D are defined in the same way. The second factor is geometric, explained in the following example:

Suppose there are only two regions: a contractile regime for $r_0 \le r < r_{23}$ and a connected regime for $r_{23} \le r < r_{12}$ (with the remainder of the domain being unconnected). The velocity profile, for the case where $\lambda_0 = \frac{2}{3}\mu_0$, is:

$$u(r) = \begin{cases} a_1 r + b_1/r, & r_0 \le r < r_{23} \\ a_2 r + b_2/r, & r_{23} \le r < r_{12} \end{cases}$$

The linear system outlined previously can be solved to yield the unique coefficient solution:

$$a_1 = -\frac{k}{2}\left(\frac{r_{12}^2 + {r_{23}}^2}{r_0^2 + r_{12}^2}\right), \qquad b_1 = +r_0^2\frac{k}{2}\left(\frac{{r_{23}}^2 + r_{12}^2}{r_0^2 + r_{12}^2}\right)$$

$$a_2 = +\frac{k}{2}\left(\frac{r_0^2 - {r_{23}}^2}{r_0^2 + r_{12}^2}\right), \qquad b_2 = +r_{12}^2\frac{k}{2}\left(\frac{r_0^2 - {r_{23}}^2}{r_0^2 + r_{12}^2}\right)$$

Interestingly, since $r_0$ is small compared to $R$, generally $b_1$ will also be small – so the velocity profile in the contractile regime is nearly linear (moreover, exactly linear as $r_0 \to 0$). Numerical solutions confirm that this is the general case in our system. Thus, $a_1$ is the contraction rate. Note that it is equal to $k$ multiplied by the geometric factor $\frac{1}{2}\left(\frac{r_{12}^2 + {r_{23}}^2}{r_0^2 + r_{12}^2}\right)$. As $r_0^2 \ll r_{12}^2$, and usually $r_{12}^2 \sim {r_{23}}^2$, this factor is on the order of unity. This turns out to be the case in general. In the numerical simulations in 2D, we use the exact expressions for the velocity profile obtained from the solutions of Eqs. 13-16 for the case where $\lambda_0 = \frac{2}{3}\mu_0$

We scale the model equations as follows. The droplet's radius, $R$, provides a natural length scale; the turnover time, $1/\beta$, is the natural time scale; and the natural density scale is $\bar{\alpha}/\beta$. We solve the model equations numerically after non-dimensionalizing the model by using these scales. In 1D, the non-dimensional model becomes (using the same notations we had for the dimensional variables for the non-dimensional ones):

$$\frac{\partial \rho}{\partial t} = \left(\varepsilon + (1-\varepsilon)\exp\left(-\rho/\rho_0\right)\right) - \rho - \frac{\partial}{\partial x}(u\rho); u = \begin{cases} 0 & \rho < \rho_{12} \\ const & \rho_{12} < \rho < \rho_{23} \\ const - kx & \rho > \rho_{23} \end{cases} \quad (17)$$

and in 2D:

$$\frac{\partial \rho}{\partial t} = \left(\varepsilon + (1-\varepsilon)\exp\left(-\rho/\rho_0\right)\right) - \rho - \frac{1}{r}\frac{\partial}{\partial r}(ru\rho); u = \begin{cases} 0 & \rho < \rho_{12} \\ (const)r + const/r & \rho_{12} < \rho < \rho_{23} \\ (const)r + const/r & \rho > \rho_{23} \end{cases} \quad (18)$$

The model has six non-dimensional parameters (for which we retain the same notations as for the dimensional parameters):

$\underbrace{\beta\rho_{12}/\bar{\alpha}}_{\text{dim}} \to \underbrace{\rho_{12}}_{\text{non-dim}}$ is the gas-connected threshold density (boundary between states I and II);

$\underbrace{\beta\rho_{23}/\bar{\alpha}}_{\text{dim}} \to \underbrace{\rho_{23}}_{\text{non-dim}}$ is he connected-contractile threshold density (between states II and III);

$\underbrace{k/\beta}_{\text{dim}} \to \underbrace{k}_{\text{non-dim}}$ (where dimensional parameter $k = \sigma_0/\eta_0$) is the contraction rate;

$\underbrace{\varepsilon/\bar{\alpha}}_{\text{dim}} \to \underbrace{\varepsilon}_{\text{non-dim}}$ is the low-density gas assembly rate;

$\underbrace{\rho_0\beta/\bar{\alpha}}_{\text{dim}} \to \underbrace{\rho_0}_{\text{non-dim}}$ is the density parameter at which the low assembly rate turns into high one;

$\underbrace{v_0/(R\beta)}_{\text{dim}} \to \underbrace{v_0}_{\text{non-dim}}$ is the effective growth rate of the percolated network boundary.

We solve the model drift-reaction equations numerically using standard numerical methods [7]. The results are relatively robust to variations of the model parameters. Specifically, we fixed several non-dimensional parameters: $\varepsilon = 0.1$, $\rho_0 = 0.5$, $\rho_{12} = 0.1$, $\rho_{23} = 0.6$. The rationale for these choices is as follows: as mentioned above, the density dependence of the assembly rate is not critical, but a reduction in assembly at low density benefits the robustness of the results: the slower assembly ensures that the contractile density is not reached prematurely before the network interconnects globally, hence the smallness of parameter $\varepsilon$ . We want the assembly rate to saturate to a constant at relatively low densities, comparable to the characteristic density scale, hence the choice of parameter $\rho_0$ . Our previous results [2] indicate that the network is interconnected at relatively low densities, so we chose parameter $\rho_{12}$ to be small. We are assuming that the network becomes contractile at densities comparable to the characteristic density scale, which means choosing parameter $\rho_{23}$ on the same order of magnitude as that of parameter $\rho_0$ . Varying the system size in the non-dimensionalized model is equivalent to changing $v_0$, which is inversely proportional to the radius. To study the behavior of the model as a function of the contraction rate and system size, we thus varied the remaining model parameters, $k$ and $v_0$. The results (Fig. 3), indicate that at small contraction rates and droplet's radii (i.e.

larger $v_0$), the density and velocity evolve to yield stable steady distributions, while at greater contraction rates and droplet's radii, periodic waves emerge. The model displays an instability at very high contraction rates ($k$ on the order of 5 and above), which are indicative of the local contractile instabilities that are discussed in detail below.

#### *Analytical estimates for the transition length and wave period*

We can make the following useful rough estimates (in 1D, but they work in 2D as well). Let us consider the case of the steady flow in a relatively small droplet of a certain radius (to be estimated), for which the free boundary of the interconnected network is exactly at the droplet's boundary. We will call the special radius at which this condition is achieved $R_{tr}$, and show below that it is equal to the transition length above which we obtain periodic waves. The network density is decreasing with distance away from the droplet's center (*x*=0). The centripetal velocity profile for this case is such that the velocity increases linearly from the *x*=0 to a distance $r_{23}$, where the density reaches the threshold contractile density: $\rho(r_{23}) = \rho_{23}$. In the region $x < r_{23}$, where the density is above the contractile threshold, the centripetal velocity profile has a slope equal to the contraction rate $k$, and so the centripetal velocity there is equal to $v = kx$ (with a negative sign, directed to the left), and the maximal value of the centripetal speed is reached at $x = r_{23}$ and is equal to $v_{\max} = kr_{23}$. At greater distances, $x > r_{23}$, the velocity plateaus: $v(x) = v_{\max} = kr_{23}$, because at such distances, the network is interconnected but does not contract.

If the free boundary of the interconnected network is stationary and positioned exactly at the droplet's boundary $x = R_{tr}$, two conditions must be satisfied at that boundary: first, the density there has to be equal to the threshold interconnected density, $\rho(R_{tr}) = \rho_{12}$. Second, the centripetal network velocity there has to exactly balance the growth rate of the network: $v_{\max} = v_0$. Thus, $v_{\max} = kr_{23} = v_0$, and as such $r_{23} = v_0 / k$. Let us now find the network density profile in the interval $r_{23} < x < R_{tr}$. The equation for the density there is: $\frac{\partial \rho}{\partial t} = \alpha - \beta\rho + v_{\max}\frac{\partial \rho}{\partial x}$. For simplicity, we will neglect the density dependence of the assembly rate and consider it constant. Also, in this outer region the density is well below the equilibrium density $\alpha / \beta$, so the disassembly term in the density equation can be approximately neglected: $\frac{\partial \rho}{\partial t} \approx \alpha + v_{\max}\frac{\partial \rho}{\partial x}$. At steady state, $\alpha + v_{\max}\frac{d\rho}{dx} = 0$, thus $\frac{d\rho}{dx} = -\frac{\alpha}{v_{\max}}$. Considering that $\rho(r_{23}) = \rho_{23}$, after integrating, we have: $\rho = \rho_{23} - \frac{\alpha}{v_{\max}}(x - r_{23})$. We use $\rho(R_{tr}) = \rho_{12}$ to obtain the following formula: $\rho_{12} = \rho_{23} - \frac{\alpha}{v_{\max}}(R_{tr} - r_{23})$. This formula then allows us to calculate the special droplet's radius $R_{tr}$:

$R_{tr} = r_{23} + \frac{v_{max}}{\alpha}\left(\rho_{23} - \rho_{12}\right)$. Considering that $v_{max} = kr_{23} = v_0$, $r_{23} = v_0 / k$, this equality can be rewritten as $R_{tr} = \frac{v_0}{k} + \frac{v_0}{\alpha}\left(\rho_{23} - \rho_{12}\right)$. Finally, based on the assumption that $\rho_{23} >> \rho_{12}$ and $\rho_{23} = c\frac{\alpha}{\beta}$, where $c$ is a dimensionless number on the order of unity, we arrive at the very useful estimate: $R_{tr} = \frac{v_0}{k} + c\frac{v_0}{\beta}$. This estimate predicts that the transition radius $R_{tr}$ is inversely proportional to the contraction rate $k$ and is proportional to the free boundary growth rate $v_0$ of the interconnected network. Fig. 3e shows that the numerical estimate of the transition radius exhibits the predicted scaling.

Let us now show that the transition length, $R_{tr}$, estimated above is the characteristic length scale separating the steady contracting flow from the periodic contraction waves. First, we show that if the droplet's radius is smaller than the transition radius, $R < R_{tr}$, then a steady centripetal flow persists. Simulations show that the smaller the droplet, the closer the contractile boundary, $r_{23}$, is to the droplet's center. This means that the maximal centripetal velocity, equal to $kr_{23}$, is smaller than $v_0$ in such droplets, and so the free interconnected network boundary is propped up against the droplet's boundary and is able to grow as fast as the centripetal retraction and to keep at the boundary. On the other hand, if the droplet's radius is greater than the transition radius, $R > R_{tr}$, then the contractile boundary moves farther away from the droplet's center. This means that the maximal centripetal velocity, equal to $kr_{23}$, is greater than $v_0$ in such droplets, and so the free interconnected network boundary moves inward. As soon as the retreating interconnected network boundary leaves a gap between its edge and the droplet's boundary, the unconnected network fragments start to appear in this gap. Their density grows, until the interconnected density is achieved, and then after a finite period of time, a new centripetal drift begins, generating periodic contraction waves. Simulations confirm this intuition (Fig S6, Supplementary Video 10).

Next, we estimate the period of contractions in droplets with radius larger than the transition size. This period is not simply equal to the time to assemble the network density at the periphery to the contractile density threshold. Indeed, let us analyze what happens after a contraction wave begins (Fig. 3d, Supplementary Video 9). First, over time $T_1$, the free network boundary retreats from the droplet's edge to a position where the interconnected network boundary transiently stops (i.e. where the centripetal contraction at the interconnected network's outer boundary is balanced by growth, so that $u + v_0 = 0$). Second, over time $T_2$, the network's density adjacent to the stopped boundary increases to the interconnected threshold. At that point, the network in the whole droplet becomes interconnected, and global contraction begins. Note that at that point, the density at the periphery has not yet reached the contractile threshold – this happens soon after global contraction begins. Thus, the period is equal to $T = T_1 + T_2$. To obtain an estimate of $T_1$, let us call the position where the network stops $r_1$, and let $w(t)$ be the position of the contracting wave's outer edge. Approximately, the centripetal contraction velocity is telescopic, and so: $\frac{dw}{dt} \approx -kw$. At the onset of the contraction, $w(t=0) = R$. The solution of this equation with such boundary condition is $w = R\exp(-kt)$, and so when the wave contracts to the position where the network stops, $r_1$, at $t = T_1$ we have the equality:

$r_1 = R\exp(-kT_1) \to T_1 = \frac{1}{k}\ln\left(\frac{R}{r_1}\right)$. Time $T_2$ is the ratio of the interconnected density threshold $\rho_{12}$ and the assembly rate $\alpha$ : $T_2 = \frac{\rho_{12}}{\alpha}$ (neglecting the small contribution from the disassembly at low density and the density dependence of the assembly rate). Note that as $\alpha / \beta$ is the equilibrium density, and $\rho_{12}$ is a fraction of this density, we can write: $\rho_{12} = C\alpha / \beta$, where $C$ is a dimensionless number. Then, $T_2 = \frac{\rho_{12}}{\alpha} = \frac{C}{\beta}$, and the time between the two consecutive waves is: $T = T_1 + T_2 = \frac{C}{\beta} + \frac{1}{k}\ln\left(\frac{R}{r_1}\right)$. Two conclusions can be made from this formula: first, the wave period is an increasing function of the droplet's radius, but the period is not very sensitive to the droplet radius because the dependence is logarithmic. Second, the period is proportional to the turnover time, $1/\beta$. Numerical simulation results confirm this intuition (Fig. S7).

To obtain a periodic wave pattern, in which the network flow is always directed inward, toward the global contraction center, the local contraction at the droplet's periphery should not initiate before a mechanical connection with the droplet's center is established. This implies that the density at the periphery should not grow above the contractile threshold during the time period between two consecutive contractions. Thus, $\frac{\rho_{23}}{\alpha} < \frac{\rho_{12}}{\alpha} + \frac{1}{k}\ln\left(\frac{R}{r_1}\right)$, so that $\frac{\rho_{23} - \rho_{12}}{\alpha} > \frac{1}{k}\ln\left(\frac{R}{r_1}\right)$.
The meaning of this inequality is simple: the left-hand-side is the time needed to grow the network from the moment it becomes interconnected to the moment it becomes contractile, whereas the right-hand-side is the time of the wave retraction. This inequality ensures that the network does not grow too fast so that it becomes contractile at the periphery before the interconnected network connects with the central core; otherwise, the contraction at the periphery becomes autonomous. Note that at a large droplet's radius this inequality becomes invalid.

If the network near the droplet's periphery reaches the contractile density threshold before the low-density network in the middle of the droplet forms a bridge to the contraction near the center of the droplet, the dynamics become more complicated. In this case, the peripheral region will begin contracting outward before eventually connecting to the central contracting core. So, the dynamics will still display global periodic contraction, but with a more complex spatiotemporal pattern. Our experimental observation under most conditions do not display such complex patterns, so we consider parameters such that the inequality $\frac{\rho_{23} - \rho_{12}}{\alpha} > \frac{1}{k}\ln\left(\frac{R}{r_1}\right)$ holds. Lastly, note that the assumption of constant assembly rates throughout the droplet is probably not realistic because recycling cytoskeletal molecules across a large droplet will become rate-limiting. Thus, for large droplets, the assembly rate at the periphery will become attenuated, and the contractile threshold there will not be reached before global interconnectedness of the network. Therefore, this makes the model we consider even more robust.

***Note about fluid vs solid network***

In the model, we assumed that the actomyosin network is a highly viscous fluid. This is well supported by multiple biophysical experiments that suggest that, on temporal scales longer than ten seconds, the actin networks behave as a fluid. The reason is that filaments' turnover and dynamic crosslinking dissipate all elastic stresses on scales of several seconds. Nevertheless, it is useful to consider the question: can a solid, elastic network explain our observations. The short answer is in quasi-1D, it can, in 2D – it cannot. Note that the network can be quasi-1D in higher dimensions, if it flows along one direction only, so the other perpendicular directions can be ignored. This is the case, for example, for a 2D cortical surface in a cylindrical geometry where the flow is directed along the cylinder's axis (as in [8]). However, the cortical network will become effectively 2D if its flow velocity depends nontrivially on both x- and y-coordinates.

In 1D, if a large segment of the network is pulled at its boundary, then this segment drifts without any deformations, regardless of whether the network is solid or fluid. Note, that the equations for a viscous fluid (assuming a certain class of network rheologies) predicts a constant velocity in space, anywhere where the contractile stress is absent, which is exactly the same as for a solid network. Therefore, whether we assume fluid or solid rheology, in a large part of the space, the network's behavior will be the same in the quasi-1D model.

The situation is dramatically different in 2D or 3D. Let us focus on 2D. If the network is viscous, the viscous stress in the axisymmetric case depends on the velocity gradient (Eq. 11) for the rheology that we assumed. The centripetal drift can be stress-free for an indefinitely long time simply if the velocity is distributed as in Eq. 14 (of course, the density could increase inward, then).

If the network is elastic, however, in the axisymmetric case, let us consider a hollow disc (annulus) geometry between $r_0$ and $R$ ( $r_0 << R$ ) . Let us denote by $U\left(r\right)$ the radial displacement of the network. Then, according to elasticity theory, the radial and tangential strains are defined as $\varepsilon_{rr} = \frac{\partial U}{\partial r}, \varepsilon_{\theta\theta} = \frac{U}{r}$ (i). The mechanical equilibrium across the network requires that the stresses satisfy: $\frac{\partial \sigma_{rr}}{\partial r} + \frac{1}{r}\left(\sigma_{rr} - \sigma_{\theta\theta}\right) = 0$ (ii). The strain-stress relations are given by Hooke's Law:

$$\varepsilon_{rr} = \frac{1+\nu}{E}\left(\left(1-\nu\right)\sigma_{rr} - \nu\sigma_{\theta\theta}\right), \varepsilon_{\theta\theta} = \frac{1+\nu}{E}\left(\left(1-\nu\right)\sigma_{\theta\theta} - \nu\sigma_{rr}\right) \quad \text{(iii)}$$

where $E$ is the Young modulus and $\nu$ is the Poisson ratio. Substituting (i) into (iii), and then the result into (ii), we arrive at the equation for the displacement: $\frac{d^2U}{dr^2} + \frac{1}{r}\frac{dU}{dr} - \frac{1}{r^2}U = 0$ (iv). This is the so-called Euler's equation, and it has the exact solution: $U = C_1 r + \frac{C_2}{r}$ (v) where $C_1$ and $C_2$ are constants. Respective formulas for the stresses are: $\sigma_{rr} = \frac{E}{1-\nu}C_1 - \frac{E}{1+\nu}C_2\frac{1}{r^2}, \sigma_{\theta\theta} = \frac{E}{1-\nu}C_1 + \frac{E}{1+\nu}C_2\frac{1}{r^2}$ (vi).

Let us now consider the following problem: we want to apply a contractile stress, $p$, to the inner boundary of the hollow disc, so that this boundary moves centripetally inward toward the central axis. Thus, $U\left(r_0\right) = -r_0$ (vii). We will use the free radial stress condition at the outer surface: $\sigma_{rr}\left(R\right) = 0$

(viii). Using (vii,viii), we can find constants $C_1$ and $C_2$. Substituting those into (v,vi) and using the condition $r_0 << R$, we can find the necessary contractile stress: $p \approx \frac{E}{1-\nu}$ and the centripetal displacement of the outer surface: $U(R) \approx \frac{2}{1-\nu}\left(\frac{r_0}{R}\right) r_0$.

The following conclusion can be made from these arguments: the contractile stress would have to be too great, on the order of the Young modulus of the solid elastic network, for even a minuscule (much smaller than the inner radius $r_0$) displacement of the outer surface. Thus, the centripetal movement of the 2D elastic solid network would be very ineffective. The centripetal drift of a multidimensional elastic network would thus require enormous contractile stresses. In order for a 2D network to contract effectively, it has to fluidize under stress. The network, of course, does not have to be a simple elasto-viscous medium to support the observed mechanics and transport in 2D; it can be a 'cable' network [9], or elasto-plastic one [10], but the point is that it should resist stretch but be easily compressed.

The effectively 1D cortical network undergoing retrograde flow observed in elongated cylindrical blebs investigated in [8] was found to be characterized by solid mechanics. This rheology benefits the hypothesized mechanism of cell propulsion through the extracellular matrix, in which a solid interconnected network assembles within the pores of the extracellular matrix allowing the cell to pull itself through the matrix. However, if the cell uses several protrusions in different direction, fluidizing the intracellular network becomes a must for the cell to move consistently; this is also essential at the cell rear. It seems that an optimal network mechanics would have a hybrid character – solid-like for extensile deformations and fluid-like for contractile deformations. Then, if the network is pulled at one end and is undergoing resulting 1D drift through a matrix or on a sticky substrate, the network behaves as a solid, maintaining its geometry and generating the necessary traction for locomotion. On the other hand, if the network must flow centripetally in 2D or 3D, then even small contractile stresses fluidize the network allowing it to contract. Note that fluidization due to the stress is sufficient for such mechanics, because the stress can simply result from internal forces (e.g., due to myosin), but the fluidization can also depend on the geometry of the system, i.e., being triggered by the convergence of the flow in 2D or 3D. To summarize, percolation and interconnectedness of the intracellular network is the common requirement for global cellular movements, while mechanical properties of the network (solid, fluid, or other rheologies) are probably adapting to specific geometry and mechanics of the environment.

***Discrete stochastic model***

The continuous model provides physical insight, yet some aspects of the pulsatile contraction are easier to address with a discrete model. To that end, we introduce a discrete stochastic model, in which instead of considering a continuous density, the network is described by a collection of points corresponding to material nodes of the network. The best ways to think about these nodes is to assume that they are focal points of microscopic contractile actomyosin units. We will only consider the 1D non-dimensional model on the interval $0 < x < 1$, the left and right ends of which correspond to the center and periphery of the droplet, respectively. The kinetics of the model involve a combination of assembly and disassembly processes. Assembly is realized by introducing a node with a constant rate $a$ at random locations. Specifically, on each small-time interval $\Delta t$, a node is introduced with probability $a \times \Delta t$. We choose $\Delta t$ small enough to keep $a \times \Delta t << 1$; the node appears at location $r$ which is a uniformly distributed random number. Disassembly occurs by having each existent node disappear with a constant rate $b$ .

The nodes interact with their two nearest neighbors to the left and to the right. The leftmost node interacts with the left boundary as if the boundary is a stable permanent node. The rightmost node does not interact with the boundary (stress-free boundary). The nearest neighbor distances between the nodes correspond roughly to the inverted density in the continuous model. Thus, the density-dependent network properties in the continuous model are replaced by interaction rules that depend on the nearest neighbor distance as follows:

1. If the inter-node distance $x$ (for each pair of the neighboring nodes) is smaller than the contractility threshold $\delta_{contr}$ ( $x < \delta_{contr}$ ) (corresponding to the interconnected and contractile high density region, $\rho > \rho_{23}$ ) then there is a contraction event. In such event, the nodes converge with a constant speed equal to $kx$ where $k$ is a constant contraction rate (on each small-time interval $\Delta t$ , $\Delta x = -kx\Delta t$ ).
2. If the inter-node distance $x$ is between the contractility threshold $\delta_{contr}$ and the connectivity threshold $\delta_{connect}$ ( $\delta_{contr} < x < \delta_{connect}$ ) (corresponding to the interconnected, but not contractile, medium density region, $\rho_{12} < \rho < \rho_{23}$ ) then the inter-node distance does not change.
3. If the inter-node distance $x$ is above the connectivity threshold $\delta_{connect}$ ( $x > \delta_{connect}$ ) (corresponding to the unconnected low density region, $\rho < \rho_{12}$ ), then the node pair does not interact.
4. At each time step, the node coordinates are $\{x_1, x_2, ..., x_{N-1}, x_N\}$ , in ascending order. We find all inter-neighbor-node distances greater than $\delta_{connect}$ ; these distances separate clusters of nodes, such that inside each cluster all neighboring node pairs interact, while neighboring clusters do not interact with each other. After the neighboring distances between the nodes in the cluster are updated according to rules 1-2, the center-of-mass of the cluster is shifted so that it does not move relative to the time moment before the interaction, which ensures frictionless telescopic actomyosin contraction of network segments with stress free boundary conditions.
5. Rule 4 works for all interacting clusters except the leftmost one. If the leftmost node of this leftmost cluster is closer to the left boundary than $\delta_{connect}$ , then after the neighboring distances are changed, the absolute positions of the nodes are shifted so that the boundary (which is a permanent node 0 in the model) does not shift. If the leftmost node of this leftmost cluster is farther from the left boundary than $\delta_{connect}$ , then we apply rule 4 for this cluster.

This discrete stochastic model is a close analogue of the continuous deterministic model. In both models, the local network behavior is separated into three distinct regimes: 1) no drift at low densities, in the unconnected regime, 2) drift without spatial convergence at medium densities, in the connected but non-contractile regime, and 3) drift with convergence at high densities, in the contractile regime. We found that in the discrete model we do not need to make the boundary of the interconnected network grow with a small constant rate to generate pulsatile contractions. The reason is that the global connectedness occurs when all nearest neighbor distances between the nodes are smaller than the threshold, which can be the case even when the innermost contracting cluster has a moving boundary, due to the discrete character of the model.

**Discrete model shows how continuous contraction, irregular pulsatile (coarsening) contraction and local contractile asters emerge:** Here we use the discrete model to demonstrate the influence of the interconnected density threshold on the character of the contractile behavior (Fig. S8). We illustrate the results of the discrete model by fixing the contraction rate $k$ , the domain size (in the non-dimensional model, the spatial domain always has the domain length equal to unity), the assembly and disassembly rates ( $a$ and $b$ , respectively), and the parameter $\delta_{contr}$ responsible for the threshold contractile density. In the simulations, we varied the parameter $\delta_{connect}$ , responsible for the threshold density at which the network becomes interconnected. The simulations shown in Fig. S8a correspond to large values of the parameter $\delta_{connect}$ . This case is equivalent to a very low interconnection threshold density, so even at low densities the network at the periphery (right) is connected to the center (left). In this case, a continuous steady contraction develops. Similar results are observed in the continuous model's simulations (Fig. 3b, Supplementary Video 9). Fig. S8b shows that as the interconnection threshold becomes closer to the contraction threshold, contractile pulses develop, starting near the periphery. When the density in the middle of the spatial domain reaches the interconnected threshold, the contraction rapidly moves the network inward, and then it takes a finite time to rebuild the density at the periphery. In this regime, which is similar to the periodic wavy contraction in the continuous model (Fig. 3c,d, Supplementary Video 9), each global contractile event starts with one or a few local contractions at the periphery, and after a characteristic random time, these contractions get connected to the continuously contracting central (left end) core of the network, the whole network globally converges to the center (left end), and the cycle repeats.

One significant drawback of the discrete model is that there is no true periodicity in such model, for a fundamental reason. When material nodes appear according to a uniform random distribution, the distances between them are distributed exponentially, so even when the average node density is low, there are always a few pairs of nodes with very small distances between them. Thus, the local contractions at random places *always* occur. These individual contractions are short-ranged, but they accumulate over time, shifting the growing interconnected clusters randomly. Ultimately, global contraction still develops over a characteristic time scale, but the contraction is not strictly periodic, because the time for bridging over the last unconnected gap in the 1D chain of nodes is random. Nevertheless, the discrete model qualitatively reproduces the continuous steady contraction and the pulsatile global contractions.

Fig. S8c shows a simulation of the discrete model for the case when the connectivity threshold is even closer to the contractility threshold. As the two thresholds get closer together, the discrete model predicts coarsening behavior of the network: local contractions merge into larger aggregates, which eventually, at irregular random time intervals and locations, connect to the droplet's center (left end of the interval) and converge there. This predicted behavior resembles the observed contractile pattern with intermediate amounts of Capping Protein or when myosin action is strengthened by Calyculin (Fig. 4). Lastly, Fig. S8d shows a simulation of the discrete model in the case when the connectivity threshold is very close to the contractility threshold. In this case, the local contractions become autonomous and never get connected to the center of the droplet (left end). These contractions randomly emerge and disappear, with clusters occasionally fusing with each other, and closely correspond to the contractile local asters observed in samples containing high concentrations of Capping Protein (Fig. 4b). Physically, these asters originate from local contractile instabilities: as soon as there is a patch of contractile density, the contraction leaves gaps at both sides of the patch. These gaps are not filled with nascent network fast enough, and so the contractions appear at random places transiently creating 'asters'.

***Possible alternative models and arguments against them***

To reiterate the principal physical problem arising from our experimental observations: the actomyosin network exhibits global periodic contraction, but given that the network does not contract everywhere all the time – how does this contraction become global, if it is initiated by local contractile events? The model we proposed above is as follows: even when the contractile network rapidly retracts from the boundary in the beginning of the periodic cycle, the assembly is still, however slow, sufficient to leave a low-density but interconnected network behind the contracting dense network. Thus, when the network reassembles to the threshold contractile density at the periphery, the peripheral contraction does not become autonomous, retracting locally to the droplet's boundary. Instead, the network becomes mechanically interconnected throughout the whole volume, making the contraction global. We favor this model because of its conceptual simplicity and because the predictions it makes explain the data well. Theoretically, there are other possible mechanisms that can interconnect local contractions and make the global contractions robust. Below, we consider such alternative possibilities and arguments against them.

***Variant of the model with global interconnected network***: One possibility is that there is not one but two interpenetrative networks – one is contractile, and another is a globally interconnected non-contractile network, with a low density, that always permeates the droplet. These networks could interact by friction and steric forces, as in theories of multiphase fluids [11]. The globally interconnected non-contractile network has mechanical properties of the so-called cable networks [9] – it can be compressed almost without resistance but has a very high resistance to stretching (it is basically a net made of ropes that can be easily coiled but cannot be stretched). Then, we expect a behavior identical to that predicted by our model. In fact, physically, the two model are very similar. Mathematically though, introducing the second network has the disadvantage of an additional complexity, which we wanted to avoid considering lack of data to discriminate between the two models.

***Model with significant friction at low network densities***: Another option for keeping the contraction global is to slow down the retraction of the network from the low-density region in the middle into two adjacent high-density contractile centers, to allow the density in the low-density region to grow faster than the outflow. In principle, this could interconnect the local contractions throughout the whole volume. The simplest mathematical way to implement this idea is to add the Darcy friction between the network and the solute in the force balance equation (in 1D): $\frac{\partial}{\partial x}\left(\eta \frac{\partial u}{\partial x}\right)+\frac{\partial \sigma_{contr}}{\partial x}=\zeta u$ . Note, that the added Darcy friction term appears on the right-hand-side. Such a term could also come from frictional interaction with the membrane/substrate in cortical networks. In the model that we explored, the friction coefficient, viscosity and contractile stress are all functions of density:

$$\eta(\rho)=\begin{cases}\eta_0, & \rho<\rho_{12}\\ \eta_1, & \rho>\rho_{12}\end{cases}, \sigma_{contr}(\rho)=\begin{cases}0, & \rho<\rho_{23}\\ \sigma_1, & \rho>\rho_{23}\end{cases}, \zeta(\rho)=\begin{cases}\zeta_0, & \rho<\rho_{12}\\ \zeta_1, & \rho>\rho_{12}\end{cases}.$$

Here $\eta_0 << \eta_1, \zeta_0 >> \zeta_1$. In other words, the network is interconnected, viscous and non-contractile for intermediate densities, between $\rho_{12}$ and $\rho_{23}$, contractile for greater densities, above $\rho_{23}$, and barely connected but experiencing great resistance from the solute at low densities, below $\rho_{12}$.
Simulations of this model showed that in small droplets there are steady distributions of density and velocity evolving, while for larger droplets, periodic waves develop (data not shown). The model works because in small droplets the absolute values of the velocity are smaller, and friction term is negligible; the network everywhere is interconnected and contracting. In large droplet, even though the new contraction starts at the periphery, the high friction at low densities prevents the contraction from depleting the low-density regions so that the reassembling network there can mechanically connects the contractile regions fast enough to make the contraction global. The problem with this model is that it is very hard to justify why the network friction would be larger at low network densities.

***Contractile ring model***: One distinct possibility suggested in [12] is that a cortex-like actomyosin shell assembles at the boundary of the droplet and when the shell becomes contractile, it shrinks centripetally like a contractile ring in dividing cells. The big difference between this and other models is that the ring model is based on circumferential, rather than radial, contractile stress. The argument against this model is that we observe waves in flat capillaries, as well as in part of the droplet with asymmetrically positioned aggregate. The observation of spiral waves also argues against the contractile ring model. To be clear, the circumferential stress does exist and contributes to the contraction; it is just not likely to be the central part of the phenomenon.

***Mechanosensitive attachment to the boundary***: Several models of the actin-adhesion clutch, supported by some experimental data, predict oscillatory periodic retrograde flow of the actin network in protrusive cell appendages [13]. The models are based on the mechanosensitive property of the actin network's adhesion to the substrate: the detachment rate is a rapidly increasing function of the pulling force. The model works as follows: there is a constant myosin-powered pulling from the rear of the protrusive cell appendage resisted initially by many firm adhesions. All adhesions are transient, and as several adhesions detach, the total myosin force per remaining adhesion increases, breaking more adhesions, and so on. As a result, an avalanche of breaking adhesions creates the so-called slipping state with rapid retrograde actin flow. Then, however, adhesions start to reattach, slowing the flow. The more the flow slows down, the smaller the force per adhesion, the more adhesions attach, eventually switching the system into the so-called gripping state, and the cycle starts again. It is possible, in principle, that such oscillations would emerge due to mechanosensitive adhesion between the network in the droplet and the droplet's boundary. The period of such oscillations would also be on the order of the network turnover time, simply because this is the characteristic time required to rebuild the retracted network, which could be the limiting step for the adhesive cycle. However, if this was the case, then the force of adhesion would be of the same order of magnitude or greater than the centering hydrodynamic force [3], and we would see pulsatile movements of the centered aggregate around the droplet. This is not the case, and so this model is unlikely.

***Diffusion-based models***: A simple possible explanation for the waves would be the following: the advection process brings the actomyosin network to the center, then the network's elements must disassemble, *diffuse to the periphery* and reassemble there for the new contraction event to start. It could be that the time to diffuse to the periphery is the limiting step determining the wave period. There are two arguments against this scenario: 1) characteristic diffusion coefficient for many cytoskeletal subunits (actin oligomers, crosslinking proteins etc) is $D \sim 1\mu m^2/s$. The characteristic diffusion time across the droplet then is $\sim R^2 / D \sim (100\mu m)^2 / 1\mu m^2 / s \sim 10^4 s$ , which is two orders of magnitude longer than the observed period of $\sim 100s$ . It is thus likely that the recycling of the cytoskeleton is local rather than global in the system. 2) wave period would increase as square of the droplet's radius, $\sim R^2 / D$ according to this model, which is not the case. This said, global diffusion of cytoskeletal molecules may have some impact on the contractile behavior, because consecutive waves of contraction sometimes have a reduced amplitude, which is likely related to insufficient diffusive transport.

There is also a more sophisticated possibility: there are many examples of periodic waves in mathematical biology models [14]. Such waves emerge when spatial diffusion is added to complex temporal dynamics that has either oscillatory or excitable, or, in general, multi-equilibrium behavior. Note that several models of actin waves on cells adhering to surfaces are based on these mathematical concepts [15]. In these cases, periodic modulation of network activity is generated by biochemical modulation of components of the actin machinery. There are two arguments against this type of model for the contractile waves: 1) There is no indication for the temporal, global complex oscillatory, excitable or multi-equilibrium behavior in the droplets (usually, complex interactions with either NPFs, or Rho GTPases, and/or adhesive complexes on the solid surface are necessary for such dynamics). Moreover, our results in droplets in which the contraction center is localized asymmetrically which show both continuous and periodic contraction further argues against the presence of global oscillatory changes in the system. 2) The wavelength in such models is $\sim \sqrt{DT}$ , where $D$ is the characteristic diffusion coefficient, and $T$ is the characteristic time scale. However, $T \sim 100s$ and $D \sim 1\mu m^2 / s$ , and so $\sqrt{DT} \sim 10\mu m$ (observed in adhering cell cortex), which is an order of magnitude smaller than what is observed in our system.

***Treadmilling polar filament array model***: Lastly, waves were predicted in a *1D mix of treadmilling actin filaments and myosin motors* [16] and observed in vitro [17]. Those are based on an intricate polarity sorting of the filaments and positive feedback between aggregation of the myosin clusters to the treadmilling actin plus ends and focusing of the plus ends in space. The characteristic length scale in this phenomenon is the filament's length, thus such phenomenon is highly unlikely on the scale of tens and hundreds of microns.

**References**


1. Kruse, K., Joanny, J.-F.c., Julicher, F., Prost, J. & Sekimoto, K. Asters, vortices, and rotating spirals in active gels of polar filaments. *Physical Review Letters* **92**, 078101 (2004).
2. Malik-Garbi, M. *et al.* Scaling behaviour in steady-state contracting actomyosin networks. *Nature Physics* **15**, 509-516 (2019).
3. Ierushalmi, N. *et al.* Centering and symmetry breaking in confined contracting actomyosin networks. *eLife* **9**, e55368 (2020).
4. Recho, P., Putelat, T. & Truskinovsky, L. Contraction-driven cell motility. *Physical Review Letters* **111**, 108102 (2013).
5. Landau, L.D. & Lifshitz, E.M. *Fluid Mechanics: Landau and Lifshitz: Course of Theoretical Physics, Volume 6*, Vol. 6. (Elsevier, 2013).
6. Alvarado, J., Sheinman, M., Sharma, A., MacKintosh, F.C. & Koenderink, G.H. Force percolation of contractile active gels. *Soft Matter* **13**, 5624-5644 (2017).
7. Garcia, A.L. *Numerical methods for physics*, Vol. 423. (Prentice Hall Englewood Cliffs, NJ, 2000).
8. García-Arcos, J.M. *et al.* Advected percolation in the actomyosin cortex drives amoeboid cell motility. *bioArxiv* (2022).
9. Paul, R., Heil, P., Spatz, J.P. & Schwarz, U.S. Propagation of mechanical stress through the actin cytoskeleton toward focal adhesions: model and experiment. *Biophysical journal* **94**, 1470-1482 (2008).
10. D'Antonio, G., Macklin, P. & Preziosi, L. An agent-based model for elasto-plastic mechanical interactions between cells, basement membrane and extracellular matrix. *Mathematical biosciences and engineering: MBE* **10**, 75 (2013).
11. Cogan, N.G. & Guy, R.D. Multiphase flow models of biogels from crawling cells to bacterial biofilms. *HFSP journal* **4**, 11-25 (2010).
12. Sakamoto, R. *et al.* Tug-of-war between actomyosin-driven antagonistic forces determines the positioning symmetry in cell-sized confinement. *Nature Communications* **11**, 1-13 (2020).
13. Chan, C.E. & Odde, D.J. Traction dynamics of filopodia on compliant substrates. *Science* **322**, 1687-1691 (2008).
14. Murray, J.D. *Mathematical biology II: spatial models and biomedical applications*, Vol. 3. (Springer New York, 2001).
15. Allard, J. & Mogilner, A. Traveling waves in actin dynamics and cell motility. *Current Opinion in Cell Biology* **25**, 107-115 (2013).
16. Oelz, D. & Mogilner, A. Actomyosin contraction, aggregation and traveling waves in a treadmilling actin array. *Physica D: Nonlinear Phenomena* **318**, 70-83 (2016).
17. Reymann, A.-C. *et al.* Actin network architecture can determine myosin motor activity. *Science* **336**, 1310-1314 (2012).

### Supplementary Figures

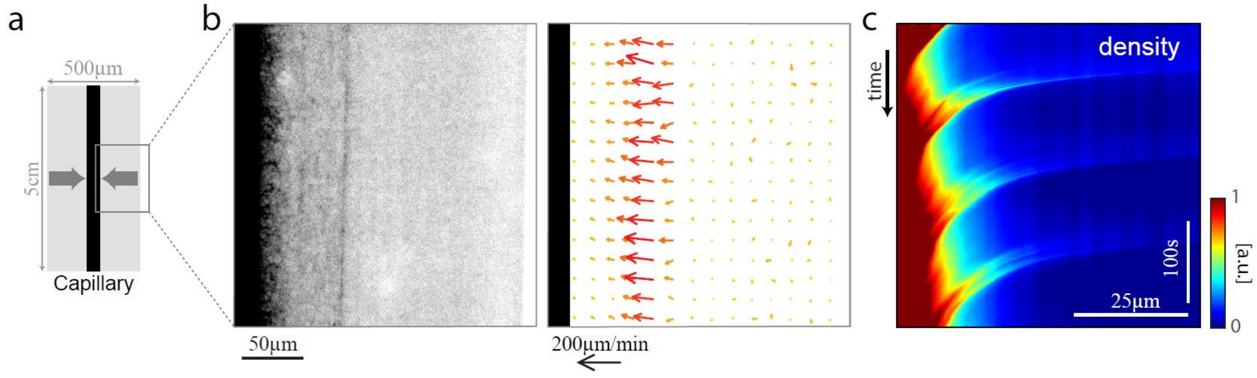


**Figure S1. Periodic contraction in an elongated capillary.** (a) Schematic illustration of contraction toward the center of a 5 cm long, 50x500 µm rectangular capillary. (b) Spinning disk confocal image (left; inverted contrast) and the corresponding velocity field (right) of a contracting wave-front parallel to the wall of the capillary, that is moving inward. The capillary was filled with 80% *Xenopus* cell extract, supplemented with lifeact-GFP to visualize the actin network. (c) Kymograph showing the density variation along a horizontal cross-section of the capillary over time. The periodic wave fronts that contract toward the center of the capillary are evident.

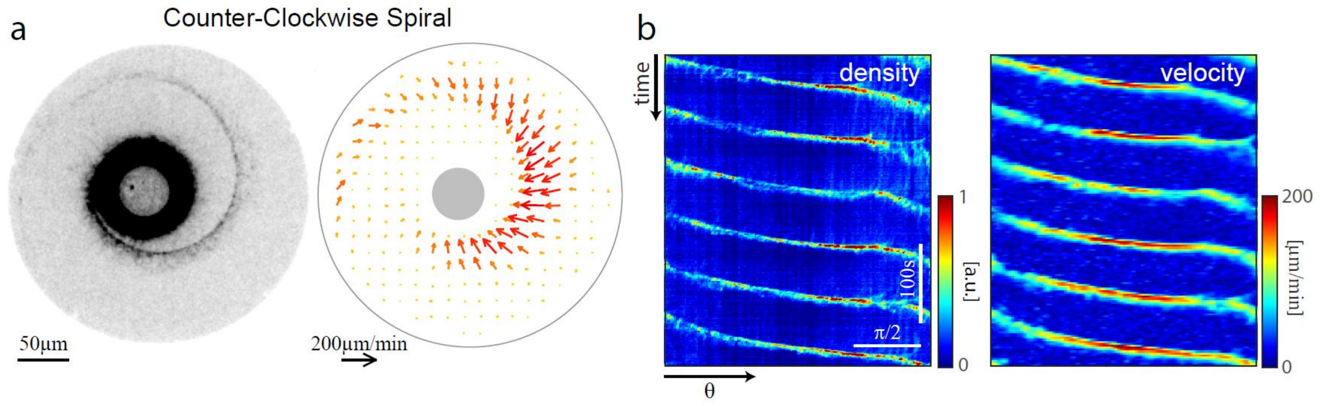


**Figure S2. Example of periodic contraction in the form of a spiral rotating counter-clockwise (CCW).** (a) Spinning disk confocal image (left) and the corresponding velocity field (right) of a water-in-oil droplet containing 98% *Xenopus* cell extract exhibiting periodic contraction in the form of a CCW spiral. (b) Kymographs showing the angular variation in the network density and velocity over time for the droplet shown in (a). The linearly-varying phase of the spiral wave front as a function of angle generates periodically-spaced parallel diagonal lines (as in Fig. 1e, but in the opposite orientation for a CCW spiral). Overall, out of 18 stable spirals observed, 10 rotated CW and 8 rotated CCW.

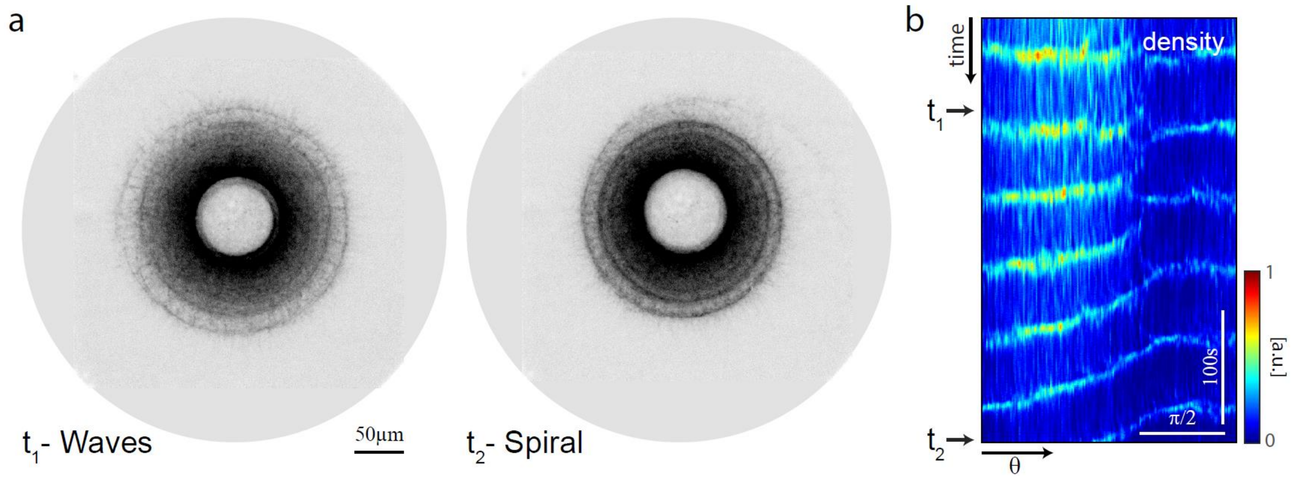


**Figure S3. Transition from concentric waves to spiral.** (a) Spinning disk confocal time-lapse images of a droplet containing 98% *Xenopus* cell extract that transitions from periodic contraction in the form of concentric waves (left) to a CW spiral (right). (b) Kymograph showing the angular variation in the network density over time for the droplet shown in (a). The transition from waves to spiral is apparent as the horizontal lines (waves) develop into diagonal lines (spiral) at a fixed angle. An angular asymmetry in the network density is present from the beginning of the movie.

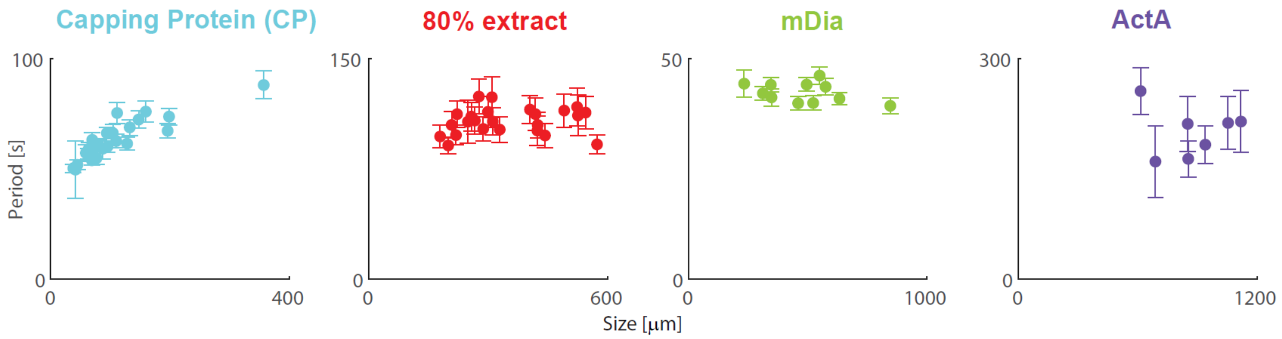


**Figure S4. Periodicity of network contraction with actin associated proteins.** Graphs depicting the wave period as a function of size for droplets above the transition length, which exhibit periodic contraction. Data is shown for populations of droplets containing 80% *Xenopus* cell extract supplemented with auxiliary proteins as indicated (1µM Capping protein, none, 0.5µM mDia, or 1.5µM ActA). The error bars indicate the uncertainty in determining the period (see Methods).

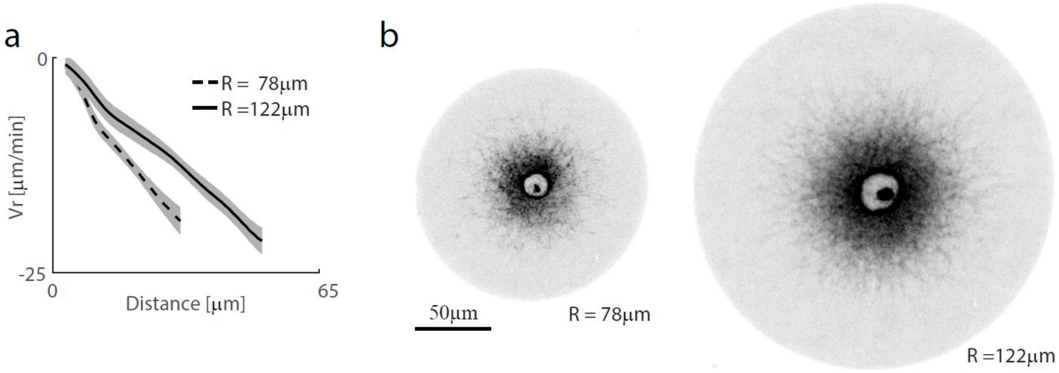


**Figure S5. Contraction rate depends on droplet size.** (a) Examples of the radial velocity profiles as a function of distance from the contraction center for two droplets of different sizes containing 80% *Xenopus* cell extract. The radial velocity increases nearly linearly in both droplets, which is a signature of homogenous density-independent contraction, but the contraction is slower in the larger droplet. (b) Spinning disk confocal images showing the actomyosin network distribution for the two droplets for which the radial velocity profiles are depicted in (a).

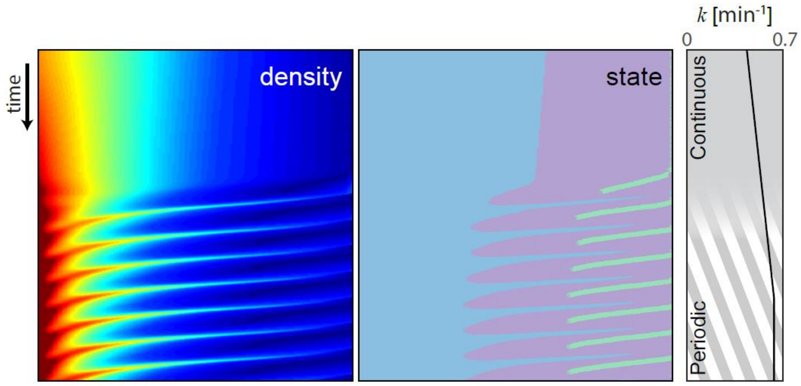


**Figure S6. Modeling the transition from continuous contraction to waves with increasing contraction rate.** Kymographs showing the evolution of the network density (left) and the corresponding local network state (center) along a radial cross-section as a function of time determined from 2D simulations of the model (Supplementary Video 10). The value of the contraction rate, *k*, was gradually increased over time from 0.45 to 0.65/min and then kept constant, as indicated in the graph on the right. The contraction dynamics transition from continuous contraction to periodic contraction.

The simulation was run taking $\rho_{12} = 0.1$, $\rho_{23} = 0.6$, and $\alpha(\rho) = 0.1 + 0.9(1 - \exp(-\rho / 0.5))$ and the following dimensional parameters: $V_0 = 30\ \mu m/min$, $\beta = 1/min$, $R = 170\ \mu m$, over a total time of 50 min.

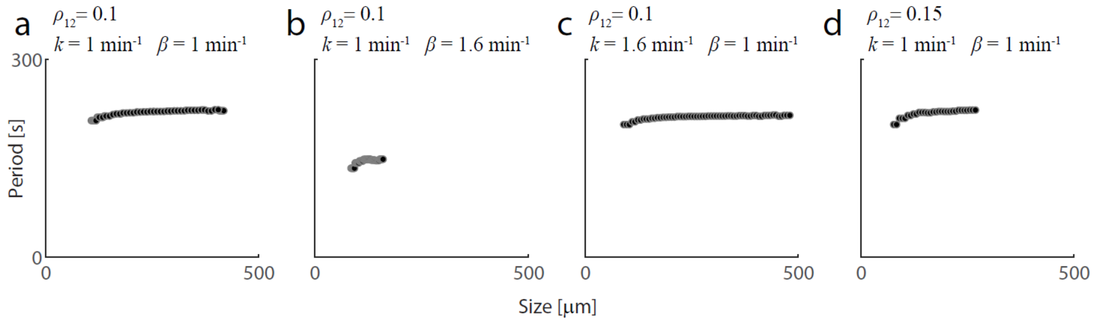


**Figure S7. Predicted wave period as a function of system size.** The period of waves in the 2D continuous model as a function of system size was determined from simulations for different values of the model parameters. The following parameters were used: $V_0 = 30$ μm/min, $r_0 = 0.1R$, $\rho_{23} = 0.6$, $\alpha(\rho) = 0.1 + 0.9(1 - \exp(-\rho/0.5))$. The other parameters varied between the four simulations as indicated in each panel. The results show that the period is not sensitive to the system size, connectivity threshold or the contraction rate, and is inversely proportional to the disassembly rate.

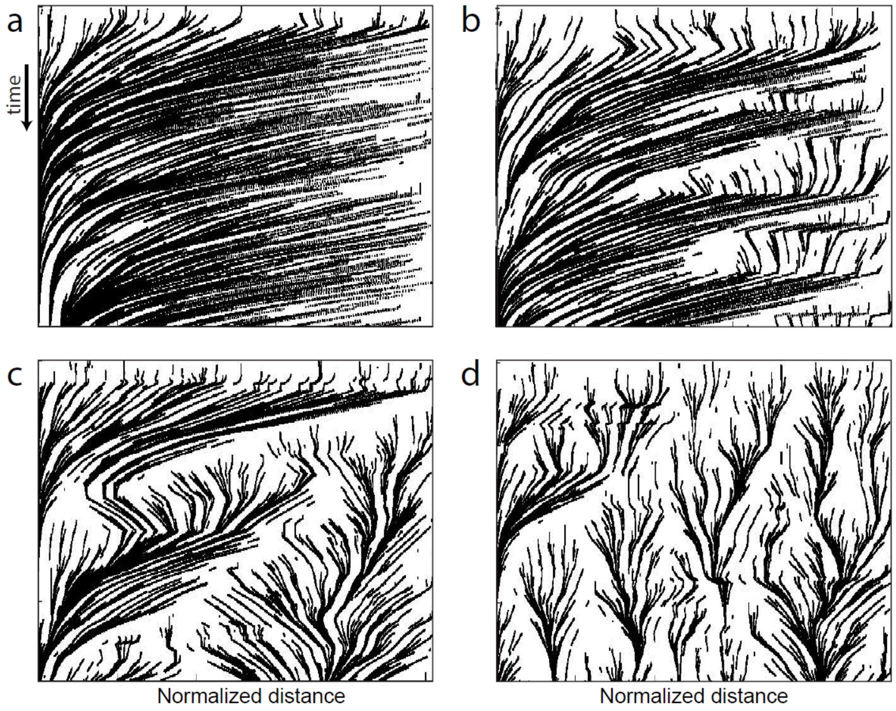


**Figure S8. Different contractile behaviors obtained from simulations of a discrete agent-based model for different parameter values.** The time evolution of the network was simulated using a discrete 1D model for different values of the connectivity threshold, $\delta_{connect}$ (see SI). Kymographs showing the location of network elements over time depict the different dynamic behaviors obtained including: (a) continuous contraction, $\delta_{connect} = 0.14$; (b) contraction waves, $\delta_{connect} = 0.08$; (c) irregular contraction, $\delta_{connect} = 0.06$, or (d) local cluster formation, $\delta_{connect} = 0.04$. All other non-dimensional model parameters were kept fixed for all simulations: contraction rate $k = 0.2$, domain size $L = 1$, assembly and disassembly rates $a = 10/$ unit time, and $b = 0.1/$ unit time, respectively (so on average there are 100 material points with an average distance of 0.01, and a turnover time equal to 10 time units), and contractility threshold $\delta_{contr} = 0.02$. The connectivity threshold, $\delta_{connect}$, was changed between simulations as indicated, and each simulation was run for a total time equal to 80 time units.

### Supplementary Videos

**Supplementary Video 1. Size-dependent contraction patterns.**

Different contraction patterns are observed in droplets of the same composition, with continuous contraction in the smaller droplet and periodic contraction in the larger droplet. This video shows spinning disc confocal images at the equatorial plane of the contracting actomyosin networks labelled with GFP-Lifeact formed within water-in-oil droplets containing 98% *Xenopus* cell extract.

**Supplementary Video 2. Continuous actomyosin network contraction.**

Steady-state continuous dynamics of an actomyosin network within a water-in-oil droplet containing 98% cell extract (Fig. 1a). This video shows spinning disc confocal images at the equatorial plane of the contracting actomyosin network labelled with GFP-Lifeact (left) and the corresponding velocity field (right).

**Supplementary Video 3. Periodic actomyosin network contraction as concentric waves.**

Periodic contraction of an actomyosin network in the form of concentric waves within a water-in-oil droplet containing 98% cell extract (Fig. 1b,c). This video shows spinning disc confocal images at the equatorial plane of the contracting actomyosin network labelled with GFP-Lifeact (left) and the corresponding velocity field (right).

**Supplementary Video 4. Periodic actomyosin network contraction in an elongated capillary.**

Periodic contraction of actomyosin network in an elongated capillary (Fig. S1). The wave-front is parallel to the boundary of the capillary and contracts towards the center of the capillary. This video shows spinning disc confocal images at the equatorial plane of the contracting actomyosin network (80% cell extract) labelled with GFP-Lifeact on one side of a 50x500 µm rectangular, 5cm long capillary.

**Supplementary Video 5. Periodic actomyosin network contraction as a clockwise spiral.**

Periodic contraction of an actomyosin network in the form of a clockwise spiral (Fig. 1d,e). This video shows spinning disc confocal images at the equatorial plane of the contracting actomyosin network (98% cell extract) labelled with GFP-Lifeact within a water-in-oil droplet (left) and the corresponding velocity field (right).

**Supplementary Video 6. Mixed contraction pattern in droplet with an off-centered contraction center.**

The dynamics of an actomyosin network in which the contraction center is positioned at an off-centered location within a water-in-oil droplet. The actin network exhibits different contractile behaviors in different regions of the droplet with periodic contraction on the far side of the droplet (upper left side) and continuous contraction on the opposite side (lower right; Fig. 1f,g). This video shows spinning disc confocal images at the equatorial plane of the contracting actomyosin network (98% cell extract) labelled with GFP-Lifeact within a water-in-oil droplet.

**Supplementary Video 7. Periodic contraction with different actin associated proteins.**

Movies of contracting actomyosin network supplemented with different actin associated proteins exhibiting periodic contraction (Fig. 2a). For each condition, spinning disc confocal images are shown at the equatorial plane of the network labelled with GFP-Lifeact. The droplets contain 80% *Xenopus* cell extract supplemented with auxiliary proteins as indicated (1µM Capping protein, none, 0.5µM mDia, or 1.5µM ActA). Note the different scale of the droplets, and the differences in periodicity.

**Supplementary Video 8. Transition from continuous to periodic contraction with increasing contraction rate.**

The dynamics of an actomyosin network in a droplet transitioning from continuous contraction to periodic contraction due to a gradual increase in the contraction rate over time (Fig. 2g,h). This video shows spinning disc confocal images at the equatorial plane of the contracting actomyosin network (98% cell extract) labelled with GFP-Lifeact within a water-in-oil droplet. The contraction pattern is continuous initially and becomes periodic.

**Supplementary Video 9. Model simulations of the size-dependent contractile behavior.**

Numerical simulations of the 2D continuous model were done for a smaller droplet (left; Fig 3b) and a larger droplet (right; Fig. 3d). The video depicts the radial density (grey) and velocity (red) profiles as a function of the normalized distance from the contraction center over time, exhibiting steady state contraction in the smaller droplet (left) and periodic contraction in the larger droplet (right; the results are shown for the non-dimensionalized variables). The colors depict regions with unconnected (green), percolated (violet) and contractile (blue) network. The simulations were run taking $\rho_{12} = 0.1$, $\rho_{23} = 0.6$, and $\alpha(\rho) = 0.1 + 0.9\left(1 - \exp\left(-\rho / 0.5\right)\right)$ and the following dimensional parameters: $V_0 = 30\ \mu\mathrm{m/min}$, $\beta = 1/\mathrm{min}$, $k = 0.8\,\mathrm{min}^{-1}$ over a total time of 30 min. The only difference between the simulations is the size of the droplet, with $\mathrm{R} = 60\ \mu\mathrm{m}$ (left) or $\mathrm{R} = 215\ \mu\mathrm{m}$ (right).

**Supplementary Video 10. Model simulations of the transition from continuous to periodic contraction with increasing contraction rate.**

Numerical simulations of the 2D continuous model were done assuming an increasing contraction rate (Fig. S6). The size of the system is initially smaller than transition length, and hence the network contracts in a continuous manner. The transition length decreases over time due to the gradual increase in the contraction rate, and the system eventually transitions into the periodic regime. The video depicts the radial density (black) and velocity (red) profiles as a function of normalized distance from the contraction center over time (the results are shown for the non-dimensionalized variables). The colors demarcate regions with unconnected (green), percolated (violet) and contractile (blue) network. The value of the contraction rate, *k*, was slowly increased over time from 0.45 to 0.65/min for the first 3/4 of the total simulation time, and then stayed constant for the last 1/4 of the time, as indicated in the graph on the right. The following parameters were used: $V_0 = 30\ \mu\mathrm{m/min}$, $\beta = 1/\mathrm{min}$, $\mathrm{R} = 170\ \mu\mathrm{m}$, $\rho_{12} = 0.1$ , $\rho_{23} = 0.6$, $r_0 = 0.1R$, $\alpha(\rho) = 0.1 + 0.9(1 - \exp(-\rho/0.5))$.

**Supplementary Video 11. Local contractions in droplets with enhanced myosin activity.**

Contracting patterns that develop in droplets containing 80% *Xenopus* cell extract supplemented with different concentration of Calyculin (Fig. 4a). This video shows spinning disc confocal images at the equatorial plane of a contracting actomyosin network (80% extract) labelled with GFP-Lifeact, supplemented with 300nM Calyculin exhibiting periodic contraction of clusters towards the center (left) or 600nM Calyculin exhibiting irregular contraction with coalescence of actin clusters (right).

**Supplementary Video 12. Local contractions in droplets with enhanced filament capping.**

Contracting patterns that develop in droplets containing 80% *Xenopus* cell extract supplemented with different concentration of Capping Protein (CP; Fig. 4b). This video shows spinning disc confocal images at the equatorial plane of a contracting actomyosin network (80% extract) labelled with GFP-Lifeact, supplemented with 1.5 µM CP exhibiting irregular global contraction of the actin network (left) and 2 µM CP exhibiting local contraction into small clusters (right).
